# RNA Polymerase II CTD phosphatase Rtr1 prevents premature transcription termination

**DOI:** 10.1101/708727

**Authors:** Melanie J. Fox, Jose F. Victorino, Whitney R. Smith-Kinnaman, Sarah A. Peck Justice, Hongyu Gao, Asha K. Boyd, Megan A. Zimmerly, Rachel R. Chan, Gerald O. Hunter, Yunlong Liu, Amber L. Mosley

## Abstract

RNA Polymerase II (RNAPII) transcription termination is regulated by the phosphorylation status of the C-terminal domain (CTD). Using disruption-compensation (DisCo) protein-protein interaction network analysis, interaction changes were observed within the termination machinery as a consequence of deletion of the serine 5 RNAPII CTD phosphatase Rtr1. Interactions between RNAPII and the cleavage factor IA (CF1A) subunit Pcf11 were reduced in *rtr1Δ,* whereas interactions with the CTD and RNA-binding termination factor Nrd1 were increased. These changes could be the result of altered interactions between the termination machinery and/or increased levels of premature termination of RNAPII. Transcriptome analysis in *rtr1Δ* cells found decreased pervasive transcription and a shift in balance of expression of sense and antisense transcripts. Globally, *rtr1Δ* leads to decreases in noncoding RNAs that are linked to the Nrd1, Nab3 and Sen1 (NNS)-dependent RNAPII termination pathway. Genome-wide analysis of RNAPII and Nrd1 occupancy suggests that loss of *RTR1* leads to increased termination at noncoding genes and increased efficiency of snRNA termination. Additionally, premature termination increases globally at protein-coding genes where NNS is recruited during early elongation. The effects of *rtr1Δ* on RNA expression levels were erased following deletion of the exosome subunit Rrp6, which works with the NNS complex to rapidly degrade terminated noncoding RNAs. Overall, these data suggest that Rtr1 restricts the NNS-dependent termination pathway in WT cells to prevent premature RNAPII termination of mRNAs and ncRNAs. Additionally, Rtr1 phosphatase activity facilitates low-level elongation of noncoding transcripts that impact the transcriptome through RNAPII interference.

**AUTHOR SUMMARY:** Many cellular RNAs including those that encode for proteins are produced by the enzyme RNA Polymerase II. In this work, we have defined a new role for the phosphatase Rtr1 in the regulation of RNA Polymerase II progression from the start of transcription to the 3’ end of the gene where the nascent RNA from protein-coding genes is typically cleaved and polyadenylated. Deletion of the gene that encodes *RTR1* leads to changes in the interactions between RNA polymerase II and the termination machinery. Rtr1 loss also causes early termination of RNA Polymerase II at many of its target gene types including protein coding genes and noncoding RNAs. Evidence suggests that the premature termination observed in *RTR1* knockout cells occurs through the termination factor and RNA binding protein Nrd1 and its binding partner Nab3. Additionally, many of the prematurely terminated noncoding RNA transcripts are degraded by the Rrp6-containing nuclear exosome, a known component of the Nrd1-Nab3 termination coupled RNA degradation pathway. These findings suggest that Rtr1 normally promotes elongation of RNA Polymerase II transcripts through preventation of Nrd1-directed termination.

## INTRODUCTION

The termination of transcription by eukaryotic RNA Polymerase II (RNAPII) is tightly coupled with RNA processing including small RNA processing, splicing, and mRNA cleavage and polyadenylation at the 3’-end of protein-coding genes [1]. Recent studies have reaffirmed that transcription termination in eukaryotes is a highly dynamic process that can lead to different gene expression outputs through mechanisms such as alternative polyadenylation site usage and premature transcription termination [2-9]. Transcription termination in yeast has been shown to be regulated through numerous termination factors as well as the phosphorylation status of the C-terminal domain (CTD) of RNAPII, which has the repetitive sequence (Tyr^1^-Ser^2^-Pro^3^-Thr^4^-Ser^5^-Pro^6^-Ser^7^)_n_ [10, 11]. However, the exact mechanisms that underlie the role of CTD dephosphorylation in the regulation of elongation, termination, and the attenuation of these processes remain unclear. At least four phosphatases are components of the yeast transcription elongation/termination machinery: Rtr1, Ssu72, Glc7, and Fcp1 [12-21]. There appears to be extensive interplay between the protein phosphatases and their control of the phosphorylation status of the RNAPII CTD. For instance, serine 5 (Ser5) dephosphorylation has been shown to be carried out by both Rtr1 and Ssu72 in both *in vivo* and *in vitro* studies [12, 22-25]. Additionally, Ssu72 dephosphorylation of Ser5 serves as a prerequisite for Ser2 dephosphorylation by Fcp1 [13, 15]. However, it remains unclear how temporal dephosphorylation impacts the formation and/or recruitment of RNA processing complexes during transcription and the determination of the termination pathway that will be used.

One pathway that is heavily influenced by CTD phosphorylation is the Nrd1, Nab3 and Sen1 (NNS) polyadenylation independent transcription termination pathway [26-31]. Nrd1 contains a RNAPII CTD interaction domain (CID) that preferentially interacts with a Ser5-P CTD [26]. The NNS complex regulates transcription termination of short non-coding transcripts and transcription elongation of a selection of protein coding genes [28, 32-35]. Pcf11, a member of the cleavage factor Ia complex (CFIa), also contains a CID that has increased affinity for Ser2-P over Ser5-P modified RNAPII CTD. Pcf11 has been shown to be required for both poly-adenylation dependent termination and NNS termination [26, 36, 37]. While the cleavage and polyadenylation factor (CPF) complex does not contain any known CID containing proteins, both Ssu72 and Glc7 are integral subunits of CPF. Rtt103 a CID-containing protein with specificity to Ser2-P CTD, has been proposed to form a higher order complex with CFIa and CPF to possibly bridge the Rat1 exoribonuclease to the RNAPII CTD to trigger degradation of the cleaved 3’ end of the RNA transcription product and hence transcription termination [26, 38]. Rat1, Rtt103, and the decapping nuclease Rai1 are sufficient to terminate elongating RNAPII *in vitro* and have been shown to be required for RNAPII termination *in vivo* [39-42]. However, numerous subunits of CFIa and CPF are required for fully efficient RNAPII termination *in vivo* suggesting that higher-order interactions between the transcription termination machinery, RNAPII, and the RNA are likely required in eukaryotes [43, 44]. Additional factors such as Rrp6, a subunit of the nuclear exosome, may also play a role in transcription termination through targeting of certain cellular states of RNAPII such as the backtracked enzyme (previously described as the reverse torpedo model) [43, 45-47]. We propose that the extensive control of RNAPII CTD dephosphorylation in eukaryotes serves as a critical regulator of co-transcriptional RNA processing and transcription termination with changes in timing of dephosphorylation by the four CTD phosphatases leading to the production of distinct transcriptional readouts.

We have previously shown that deletion of the Ser5-P CTD phosphatase Rtr1 results in an increase in global Ser5 RNAPII phosphorylation and disruption of termination at specific protein-coding genes [12, 48]. In this current work, data shows that *RTR1* deletion leads to alterations in CID-containing protein interactions with the RNAPII CTD. Additionally, the interactions between CFIa and CPF are decreased in the absence of Rtr1, suggesting that the timing of CTD dephosphorylation may regulate the formation of stable interactions between the termination machinery, the nascent RNA, and RNAPII. Transcriptome analysis reveals that *rtr1Δ* cells have decreased levels of a variety of noncoding transcripts. Analysis of Nrd1 occupancy shows that both Nrd1 and RNAPII accumulate at known Nrd1 binding sites in WT cells but that *RTR1* deletion causes a decrease in RNAPII and Nrd1 levels at both noncoding and coding genes, suggesting increased elongating RNAPII turnover through termination. This study shows that Rtr1 is globally required for the prevention of premature termination of RNAPII at protein-coding genes by reducing the efficiency of termination through the NNS-dependent termination pathway. Since the NNS termination pathway is known to recruit the RNA exosome to carry out termination-coupled RNA processing and/or decay, the impact of *rtr1Δ rrp6Δ* was also explored, revealing that the decreases in noncoding RNA levels in *rtr1Δ* require Rrp6 activity. These findings clearly show that Rtr1 is required to attenuate termination through the NNS pathway. Overall, our findings suggest that precise control of CTD dephosphorylation is required to maintain the balance between elongation and termination at the wide variety of target genes whose transcripts are produced and co-transcriptionally processed by RNAPII.

## RESULTS

### Disruption-compensation analysis reveals changes in termination factor interactions in *RTR1* deletion cells

The phosphorylation status of the RNAPII CTD plays a major role in the regulation of the mechanisms through which transcription termination occurs [49-52]. We have recently shown that deletion of *RTR1* causes global increases in CTD Ser5-P [48] and it has previously been shown that loss of *RTR1* results in 3’-end processing defects at the polyA-dependent gene *NRD1* [12]. To determine the role of Rtr1 in the regulation of RNAPII interactions with termination factors, we performed <u>Dis</u>ruption-<u>Co</u>mpensation (DisCo) network analysis. It has been postulated that genetic perturbations can cause edge-specific changes in a protein-protein interaction (PPI) networks. DisCo combines genetic perturbation and in-depth affinity purification-mass spectrometry (AP-MS) studies to obtain unique biological insights into the mechanisms that cause phenotypic changes in gene expression networks. For these studies, we generated dynamic protein-protein interaction networks using Significance Analysis of INTeractome (SAINT) probability scores in the presence or absence of Rtr1 [53, 54]. At least four biological replicate affinity purifications were performed for Nrd1-TAP, Pcf11-TAP, and Ssu72-FLAG to represent the Nrd1-Nab3, cleavage factor Ia (CFIa), and cleavage and polyadenylation factor (CPF) complexes, respectively, from either WT or *rtr1Δ* cells. The resulting data matrix consists of 24 x 3,960 protein-level measurements in 668,969 peptide-spectrum matches (PSMs, Table S1). We focused our analysis on high-confidence interactions between the protein components of the termination factor complexes along with the two largest subunits of RNAPII, Rpb1 and Rpb2 although a lower stringency network prepared using STRING v11 is also included (Figure S1, [55]).

Prey-prey correlation analysis was performed for all purifications from WT or *RTR1* deletion genotypes. In brief, a high correlation value between two proteins suggests that they have a similar distribution of PSMs across the same set of purifications independent of the bait protein used for purification. Proteins that function within the same protein complex typically have the highest correlation values, as shown in Fig. 1A. In addition, there is evidence of cross-association of cleavage factor Ia (CF1a), the cleavage and polyadenylation factor (CPF) and RNAPII (Pol II) in WT cells through a positive association between CFIa and the CPF subunits Fip1 and Pap1, which are both components of the recently described poly(A) polymerase module of CPF (Fig. 1A, [56]). Analysis through SAINT probability calculation revealed association of additional CPF subunits with CFIa (Fig. 1C). These data support a previous model which suggested that the CTD-interaction domain (CID) of Pcf11 facilitates formation of a CPF-CF1a-RNAPII complex for stable 3’-end complex formation and mRNA polyadenylation [26]. However, in cells lacking the CTD phosphatase Rtr1, the cross-correlation between the CFIa and CPF complexes is markedly reduced (Fig. 1B). In addition, the correlation between CFIa and RNAPII is also reduced, suggesting that deletion of *RTR1* leads to reduced interactions between Pcf11, the bait protein for CFIa, and RNAPII.

**Figure 1:**
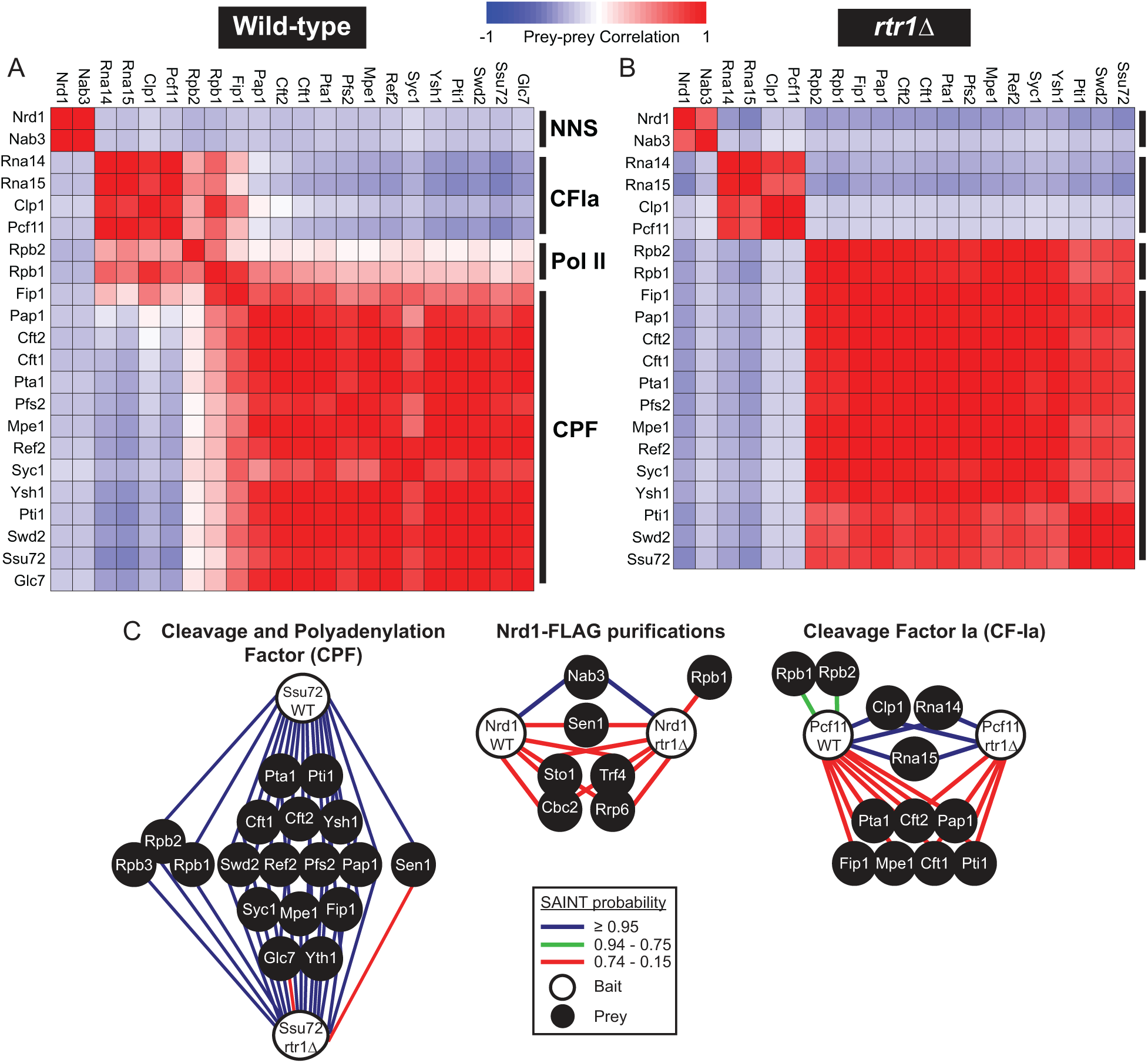
DisCo network analysis of complexes involved in RNAPII transcription termination in WT and *rtr1Δ* yeast. A) Prey-prey correlation analysis of yeast termination complex affinity purification-mass spectrometry data from WT cells of ≥ 4 biological replicate purifications for each bait protein (Nrd1, Pcf11, and Ssu72, total n=12). B) Prey-prey correlation analysis of yeast termination complex affinity purification-mass spectrometry data from *rtr1Δ* cells of ≥ 4 biological replicate purifications for each bait protein (Nrd1, Pcf11, and Ssu72, total n=12). C) Ssu72-FLAG, Pcf11-TAP and Pcf11-FLAG, and Nrd1-TAP protein-protein interaction networks from WT and *rtr1Δ* cells depicting SAINT analysis of ≥ 4 biological replicate purifications for each genotype. The nodes represent a protein of interest whereas the edges represent the SAINT interaction probability as indicated in the legend. Each protein-protein interaction network represents a subset of the full protein-protein interaction network which is shown in the supplement with a fold-change cutoff value of 5 for each WT purification (Figure S1).

SAINT analysis provides additional insights into the interactions between the termination machinery through interaction probability calculation. Ssu72 is a known member of the CPF complex [13], and all CPF subunits had SAINT probabilities of ≥ 0.95 (Fig. 1C, Table S2). When isolating protein complexes through affinity purification-mass spectrometry, we have found that protein complex subunits typically have SAINT probabilities in the 0.95-1 range as we have observed with Ssu72-FLAG and the subunits of CPF. However, proteins that interact with the protein complex of interest can display a wide-range of SAINT probabilities that likely reflect the dynamic nature of some protein-protein interactions. Of note, dynamic interaction partners of RNAPII that regulate different stages of transcription were previously assigned SAINT probability values that ranged from 0.23 (Spt16) to 1 (Spt5, Pta1, Tfg1, Tfg2) in replicate Rpb3-TAP purifications [57]. In the Ssu72-FLAG purifications, the two largest subunits of RNAPII were detected as significant Ssu72 interacting proteins in both WT and *rtr1Δ* cells (Fig. 1C). Rpb1 and Rpb2 have the highest number of detectable peptides amongst the twelve RNAPII subunits and therefore the highest probability of detection in an affinity purification-mass spectrometry approach [58]. Altogether, these data suggest that loss of Rtr1 function does not alter the interactions between the Ssu72-copurifying CPF complex and RNAPII, in agreement with the prey-prey correlation analysis (Fig. 1).

CFIa has been characterized as a four-subunit protein complex containing Rna14, Rna15, Clp1, and the CID-containing protein Pcf11. SAINT analysis of biological replicates of Pcf11-TAP and Pcf11-FLAG purifications from WT and *rtr1Δ* cells (n ≥ 4) identified these four CFIa subunits with probabilities ≥ 0.95, supporting their designation as a protein complex (Fig. 1C, Table S2). Previous studies have clearly shown that the purified Pcf11 CID domain has lower affinity for Ser5-P and Ser2,5-P than Ser2-P modified CTD [26]. In *RTR1* deletion cells, we have shown that Ser2,5-P CTD repeats are present further downstream than in WT cells, as supported by increased histone H3K36me3 levels in *rtr1Δ* cells [48]. As illustrated in Fig. 1C, we find that the RNAPII subunits Rpb1 and Rpb2 interact with Pcf11 in WT cells with a SAINT probability of 0.75 (Table S2). However, no statistically significant interaction was detected between Pcf11 and Rpb1/Rpb2 when isolated from cells lacking Rtr1, as also observed from prey-prey correlation analysis (Fig. 1B & C, Table S2). It is possible that the interaction between Pcf11 and Rpb1 still occurs in *RTR1* deletion cells, but that it was below the limit of detection for our affinity purification-mass spectrometry studies. Of interest, a number of CPF subunits were found in single-affinity step Pcf11-FLAG purifications at low levels, which are more apparent with SAINT analysis than through correlation analysis. Interestingly, the SAINT probabilities for CPF subunit interactions with CFIa were similar in WT and *RTR1* deletion cells although fewer subunits of CPF were recovered (Fig. 1C). These findings suggest that the interactions between the CFIa and CPF complexes occur in the absence of stable Pcf11-CTD interactions that were not detected in *RTR1* deletion cells. It is likely, however, that CFIa-CPF interactions are strengthened through association with the nascent RNA and are further stabilized through interactions with the RNAPII CTD.

Nrd1 has been proposed to function within a protein complex containing Nab3, Sen1, Cbc2, and Sto1 [29]. However, in our quantitative proteomics analysis of Nrd1 affinity purifications, only Nab3 was found to interact with Nrd1 with a SAINT probability of ≥ 0.95, and this interaction was also the only high correlation prey-prey value found (Fig. 1). Sen1, Sto1, and Cbc2 were identified as Nrd1 interacting proteins although their SAINT probability values indicate they are dynamic interacting partners of the bait, Nrd1 (Fig. 1C). Additionally, subunits of the TRAMP complex and the nuclear exosome were also identified as dynamic interacting partners of Nrd1 consistent with previous findings (Fig. 1C, [29]). The Nrd1 CID has been shown to have the highest affinity for Ser5-P and Ser2,5-P CTD repeats [50], whose abundance is increased in *RTR1* deletion mutants [12, 48]. As illustrated in Figure 1C, an interaction between Nrd1 and Rpb1 was detected with a SAINT probability of 0.49 in *RTR1* deletion cells (Table S2). These findings suggest that loss of *RTR1* increases the interaction between the Nrd1-Nab3 complex and RNAPII *in vivo*, likely due to increases in the number of Ser5-P modified CTD repeats. Even in *rtr1Δ* cells the interaction probability between Nrd1 and Rpb1 was lower than what was measured for Pcf11 from WT cells. This may suggest that the Nrd1-Rpb1 interaction occurs at a lower frequency than the Pcf11-Rpb1 interaction, which is consistent with previously measured binding affinities for each CID.

### Rtr1 regulates noncoding RNA expression

To determine how global RNAPII transcription was altered upon loss of the RNAPII CTD phosphatase Rtr1, we performed strand-specific RNA-Seq analysis of total RNA from four biological replicate RNA purifications. ERCC spike-in control was included to detect the presence of global transcription defects [59]. Following alignment, differentially expressed transcripts were identified using edgeR analysis (Fig. 2, Table S3, [60]). Previously defined transcript annotations were used to distinguish multiple types of RNAPII transcripts including small nuclear/nucleolar RNAs (sn/snoRNAs), open reading frame transcripts (ORF-Ts), cryptic unstable transcripts (CUTs), stable unannotated transcripts (SUTs), and Nrd1-unterminated transcripts (NUTs) [61, 62]. Transcripts that are antisense to ORF-Ts were annotated as antisense transcripts (ASTs) [63]. In total, there was a reduction in 1481 transcripts in *RTR1* deletion cells including a large number of ASTs and other ncRNA transcripts (Fig. 2A & B). Two-hundred and seventy-six transcripts showed upregulation of more than 1.5-fold, many of which were ORF regions. The most significantly reduced transcript was *IMD2*, a well-described target of the NNS termination pathway whose expression is regulated by an intergenic NNS terminator (Fig. 2A, labeled in green, [34, 35, 64, 65]). The transcript for *NRD1* was also significantly reduced in *rtr1Δ (*Fig. 2A, labeled in green)*. NRD1* is known to be regulated by premature RNAPII termination through the Nrd1-Nab3 pathway as a form of auto-regulation [32]. Overall, these data suggest that loss of Rtr1 activity results in the downregulation of a number of different classes of ncRNAs with additional changes in specific mRNAs. These data suggest that the increased Ser5-P RNAPII CTD levels in *rtr1Δ* cells causes elevated activity of the NNS-dependent termination pathway, stimulating premature transcription termination.

**Figure 2:**
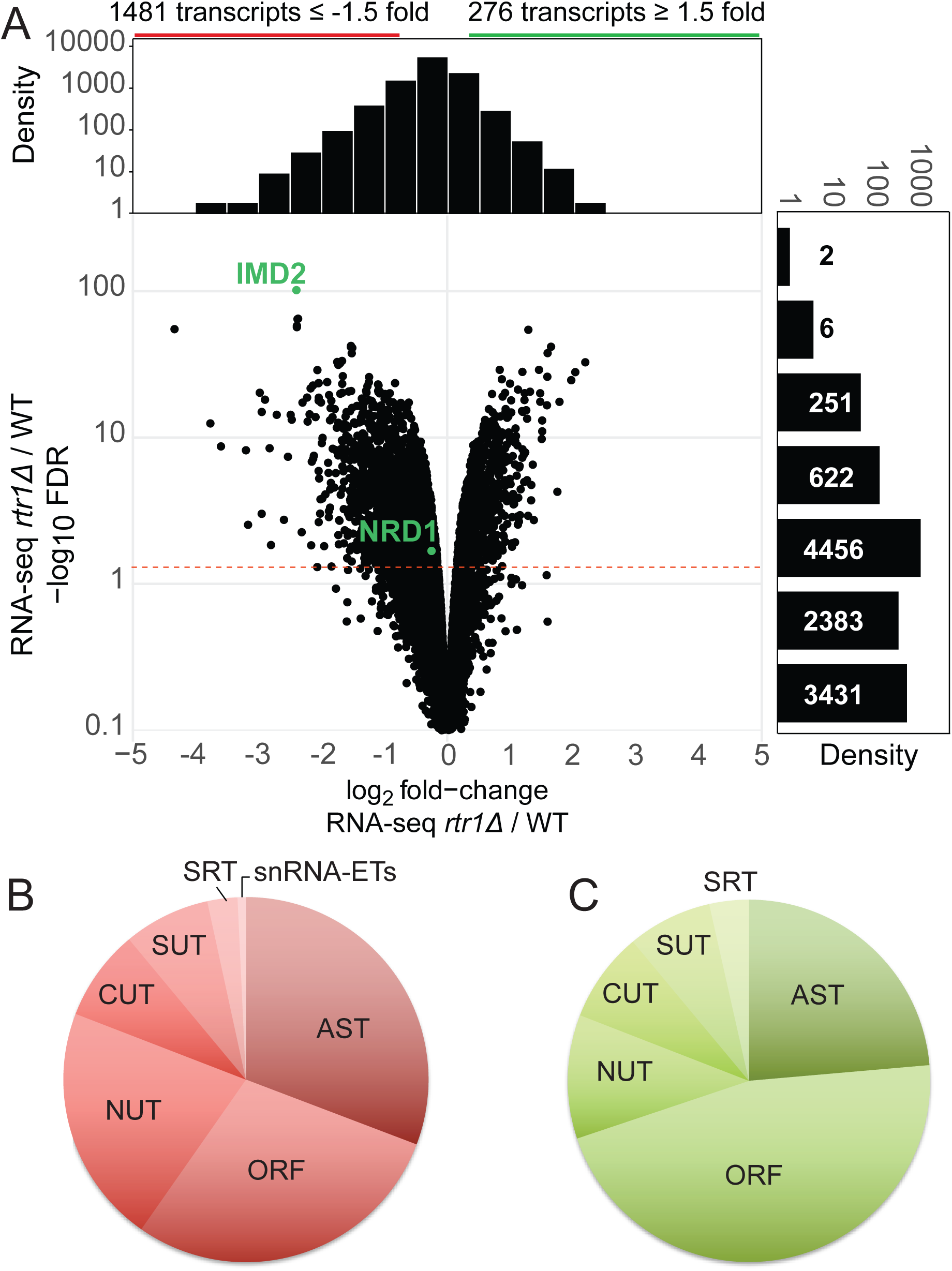
Loss of Rtr1 activity causes wide-spread changes in noncoding RNA expression in yeast. A) Volcano plot for normalized RNA-Seq data for noncoding transcripts in *rtr1Δ* vs. WT cells (n=4, edgeR analysis). Data is shown for all annotated CUTs, NUTs, and ASTs. FDR cutoff of ≤ 0.05 is indicated with a dashed red line. Density plots are shown relative to each axis for comparison. B&C) Distribution of transcripts differentially expressed in *rtr1Δ* cells compared to WT across annotation categories. Downregulated transcripts are represented in the red diagram while upregulated transcripts are represented in the green diagram.

RNAPII and Nrd1 occupancy was measured genome-wide through chromatin immunoprecipitation followed by exonuclease digestion and genome-wide sequencing (ChIP-exo) as described previously [66]. Considering that Nrd1 does not bind DNA directly, rather it binds nascent RNA and RNAPII CTD repeats at Ser5-P; we predicted that ChIP-exo of Nrd1-TAP would detect regions of DNA bound by RNAPII in complex with Nrd1 (Fig. 3A). To confirm that the binding patterns observed were specific to Nrd1, we compared the Nrd1-TAP ChIP-exo normalized read counts from WT cells to those of Rpb3-FLAG (Fig. 3B). *URA8* is known to be regulated by alternative start site selection that is dependent on nucleotide availability and is a known target for NNS-dependent early termination [30, 62]. The *SOD1* locus is convergent with *URA8* and lacks RNA binding sites for Nrd1. Figure 3B illustrates the differences seen in the binding patterns of total RNAPII (Rpb3-FLAG) and Nrd1-bound RNAPII (Nrd1-TAP) when comparing transcripts with high (*URA8*) and low (*SOD1*) levels of consensus Nrd1-Nab3 RNA binding sites. The consensus Nrd1 binding site of TT<u>TGTAA</u>AGTT is located 40 nt upstream of the *URA8* ATG. The alternative start site is terminated by the NNS pathway in nutrient-rich conditions such as growth in YPD as used in this study. Our ChIP-exo analysis of Rpb3-FLAG shows that RNAPII is localized at the 5’-end of the *URA8* gene and throughout the *SOD1* coding region (Fig. 3B). The 5’-end localization of RNAPII at URA8 corresponds with the peak of Nrd1 binding in the same area, supporting previous work that found that the majority of *URA8* transcript is terminated in early elongation by the NNS pathway, resulting in low-level transcription of full-length *URA8* [67]. The levels of Nrd1 association at the *SOD1* gene are much lower than at *URA8* even though total Rpb3-FLAG occupancy is relatively higher at *SOD1* than at *URA8* confirming that we are able to obtain selective enrichment of Nrd1 on chromatin using the ChIP-exo approach.

**Figure 3:**
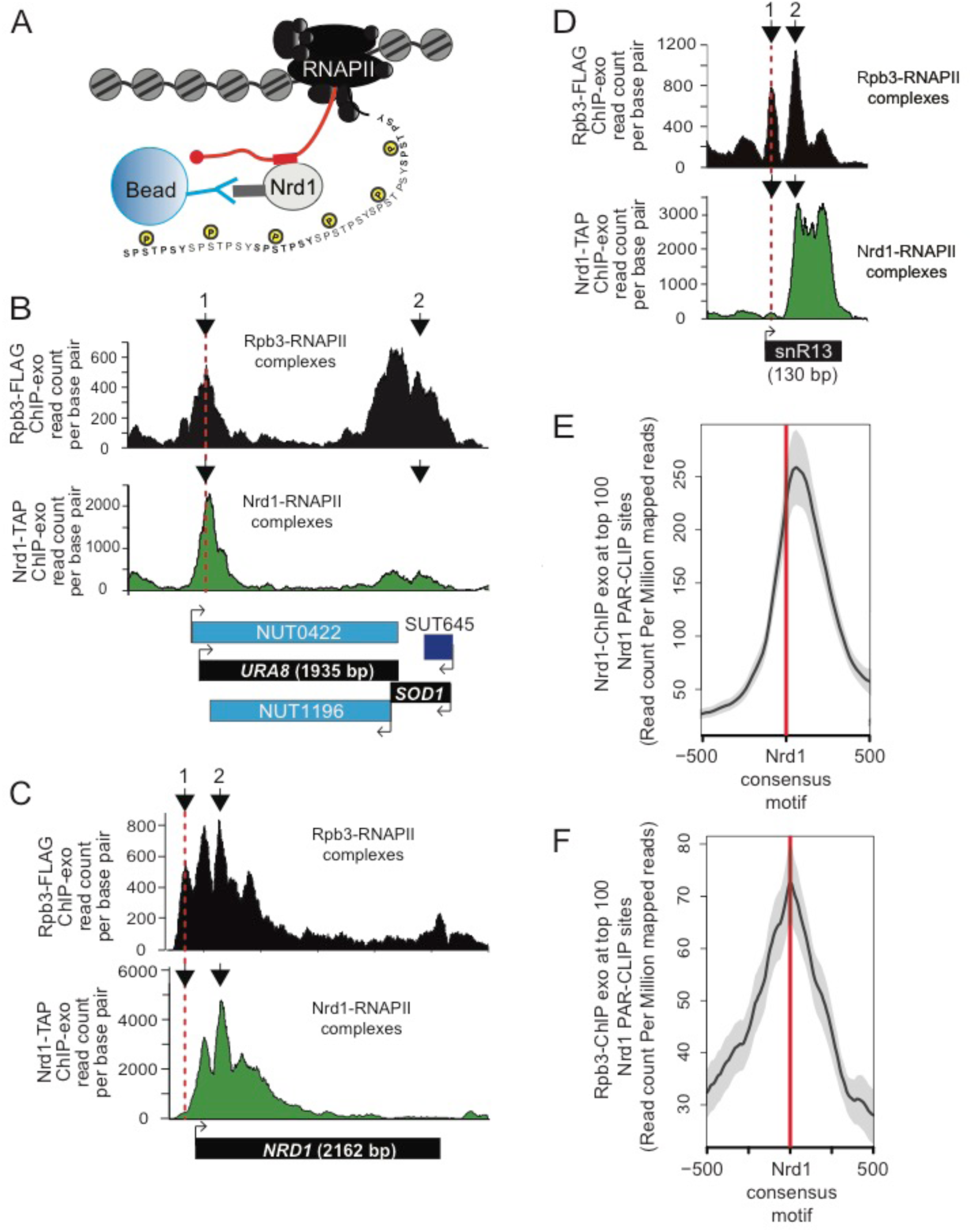
Analysis of RNAPII and Nrd1 occupancy throughout the genome in WT and *rtr1Δ* cells. A) Simplified schematic of immunoprecipitation of Nrd1-TAP by IgG-sepharose beads. Nrd1 binds to sequences in the RNA (red rectangle) via its <u>R</u>NA <u>R</u>ecognition <u>M</u>otif (RRM domain). The RNA consensus sequences for Nrd1 binding are UGUAG and UGUAA. Nrd1 binds RNAPII at Ser5-P of the CTD of RNAPII via the Nrd1 CID. Therefore, DNA sequences detected by Nrd1-TAP ChIP represent regions of DNA bound by RNAPII bound to Nrd1. B) Graphical representation of Rpb3-FLAG (black) and Nrd1-TAP (green) occupancy in the WT strain as determined by ChIP-exo sequencing reads mapped to *URA8-SOD1*. Data is shown from a representative biological replicate. Points of interest are noted by arrows. C) Occupancy of RNAPII and Nrd1 at the *NRD1* gene by ChIP-exo. Points of interest are noted by arrows. D) Occupancy of RNAPII and Nrd1 at the *SNR13* gene by ChIP-exo. Points of interest are noted by arrows. E) Average gene analysis of Nrd1 occupancy at the top 100 most enriched Nrd1 RNA binding locations throughout the yeast genome. F) Average gene analysis of RNAPII (Rpb3) occupancy at the top 100 most enriched Nrd1 RNA binding locations throughout the yeast genome.

Upon analysis of the Nrd1 ChIP-exo dataset, a pronounced peak of RNAPII was observed upstream of the location of well-positioned Nrd1-RNAPII complexes (Fig. 3C, arrow 1 vs. arrow 2). These peaks were observed at both protein-coding gene NNS targets such as *NRD1* (Fig. 3C) and noncoding genes such as *SNR13* (Fig. 3D, compare arrow 1 for upstream peak to arrow 2 for Nrd1-RNAPII peak). This suggests that Nrd1 binding to RNAPII may produce a stably paused RNAPII at strong NNS consensus sites. To further explore these findings, we used previously published Nrd1 PAR-CLIP datasets to annotate the top 100 most intense sites and then averaged the Nrd1 and RNAPII intensities surrounding the Nrd1 consensus RNA binding sites [30, 31]. In Fig. 3E, we observed a narrow peak of Nrd1-RNAPII complexes located just downstream of the genomic location of the Nrd1 consensus motif (marked with a red line). These data suggest that the pausing of RNAPII occurs just downstream of the genomic location of the Nrd1 consensus sequence, perhaps as the cognate RNA binding motif extends outside the RNA exit channel of RNAPII. A similar peak is observed from the Rpb3 ChIP-exo, although the peak is somewhat 5’ shifted, perhaps as an average of the Nrd1 bound and unbound RNAPII populations (Fig. 3F). Globally, Nrd1 was also found to localize to the 5’-end of most mRNA encoding genes and, in agreement with previous studies using ChIP-microarray analysis, the mRNA peak of Nrd1 occupancy occurs 93 +/- 3 nucleotides downstream of the annotated mRNA transcription start sites (TSS, Figure S2).

### RNA Polymerase II and Nrd1 occupancy are reduced at snRNA genes in *rtr1Δ*

Total RNA-Seq analysis revealed changes in a number of ncRNA classes including snRNAs (Fig. 2). This includes snRNA 3’-ends that we have manually annotated as extended transcripts (ETs), which are the regions downstream of snRNAs that are within the zone of termination [1]. Full snRNA transcripts are subsequently subjected to 3’-end processing through the NNS-termination pathway in coordination with the TRAMP complex and the Rrp6-containing RNA exosome [1, 29, 68-72]. Average gene analysis was performed using the ChIP-exo datasets for Rpb3 and Nrd1 for the snRNA genes aligned to the TSS with 500 bp of data upstream and 1kb downstream [73]. As shown in Fig. 4A, the average RNAPII signal at snRNA genes is decreased in *rtr1Δ* cells relative to WT, as is the Nrd1 occupancy. The relative enrichment of Nrd1 to RNAPII is also decreased at snRNA genes, perhaps suggesting that less Nrd1 recruitment is required to mediate termination in cells lacking Rtr1. In fact, the overall reduction in RNAPII levels at snRNAs suggests that termination may occur in a more rapid fashion, thus causing reduced overall RNAPII levels. In support of this hypothesis, the abundance of extended snRNA transcripts is also decreased in *rtr1Δ* cells in multiple cases, suggesting that termination occurs earlier. If overall snRNA transcription were reduced, we would also expect to measure a decrease in the mature snRNA levels. However, the overall levels of the mature snRNAs for snR62, 32, 189, and 46 are not statistically different than WT, suggesting that the major changes in these RNA occur at their 3’-ends.

**Figure 4:**
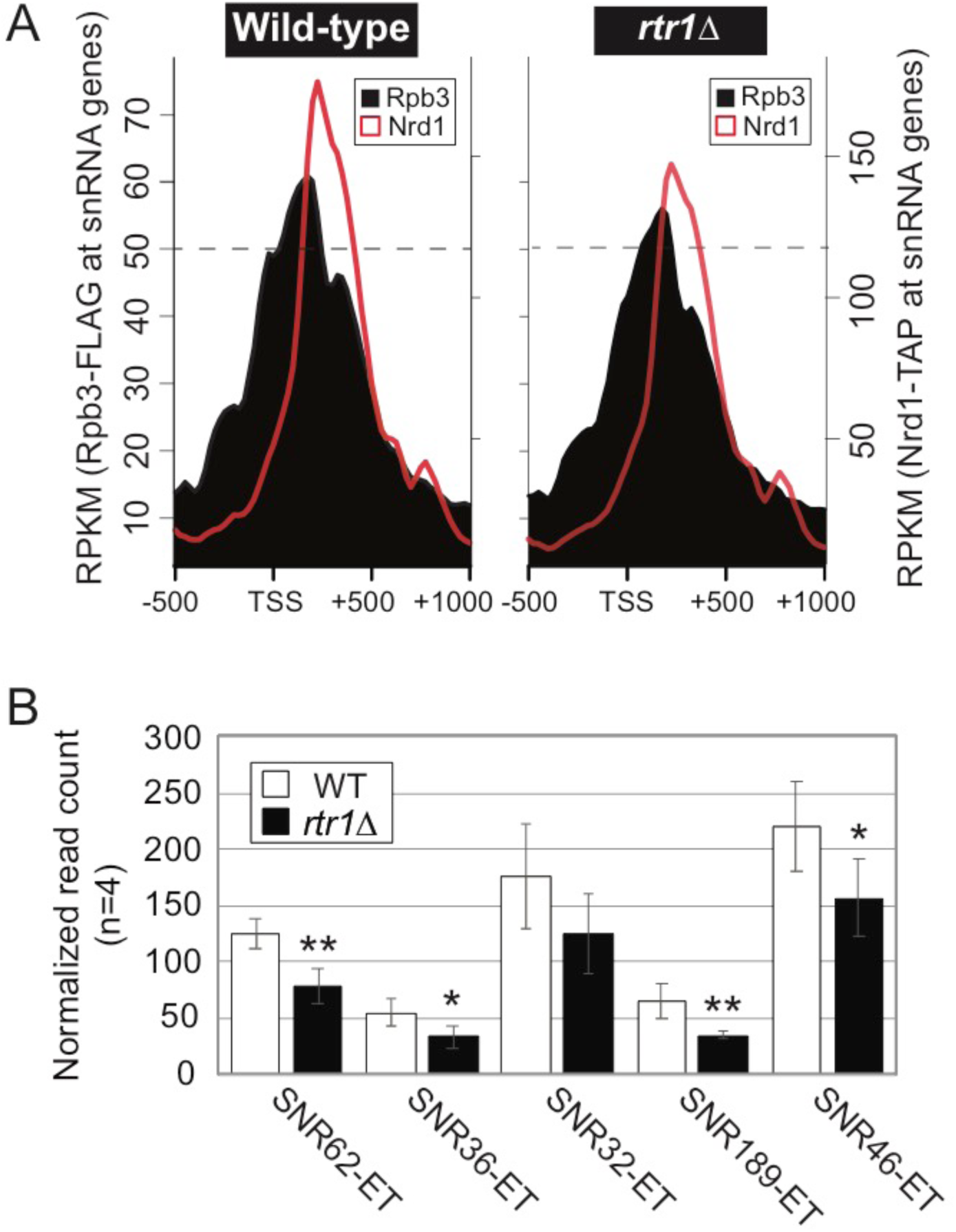
Loss of Rtr1 activity leads to decreased RNAPII occupancy at snRNA genes and shortened snRNA transcripts. A) Average gene analysis of RNAPII and Nrd1 occupancy at snRNA genes in WT (left) or rtr1Δ (right) cells. The data is shown as average RPKM values calculated using ngs.plot −500 and +1000 nucleotides relative to the snRNA gene annotated TSSs in either WT or rtr1Δ cells. B) RNA-Seq read count values for manually annotated extended transcript regions from a subset of snRNA genes.

### Global levels of RNA Polymerase II and Nrd1 occupancy are altered in *RTR1* deletion cells

Although Nrd1 recruitment is highest at RNAPII target genes containing RNA binding sites for Nrd1-Nab3 (such as *URA8*), average gene analysis in this study and others has shown that Nrd1 is recruited just downstream of the peak of RNAPII Ser5-P CTD phosphorylation at protein-coding genes [30, 37]. At the model protein-coding gene *PMA1*, RNAPII occupancy is relatively consistent across the entire length of the gene (data not shown). Nrd1 binding, in contrast, peaks in the 5’ end of the gene, ∼270-321 nt past the annotated transcription start site of *PMA1.* To measure changes that occur in overall RNAPII occupancy in cells +/- Rtr1 we compared the localization of Nrd1-TAP and Rpb3-FLAG at protein-coding genes 1000 nucleotides downstream of the TSS and compared this to histone occupancy data we obtained by MNase-Seq from WT cells (Fig. 5). The overall RNAPII and Nrd1 occupancy observed relative to the annotated transcript end site (TES) +/- 200 nucleotides was also determined. Rpb3 localization is slightly higher at the 5’ end of protein-coding genes in *RTR1* deletion cells relative to WT. However, Rpb3 occupancy decreases in *rtr1Δ* more than in WT cells as RNAPII progresses towards the 3’-end of these genes and at the TES (Fig. 5A). Nrd1 levels show a small but consistent decrease in *rtr1Δ* samples relative to WT across the entire average gene and at the TES. Interestingly, the transition to lower levels of RNAPII occupancy in *rtr1Δ* relative to WT occurs just following the peak of Nrd1 recruitment (overlaid data shown in Figure S3).

**Figure 5:**
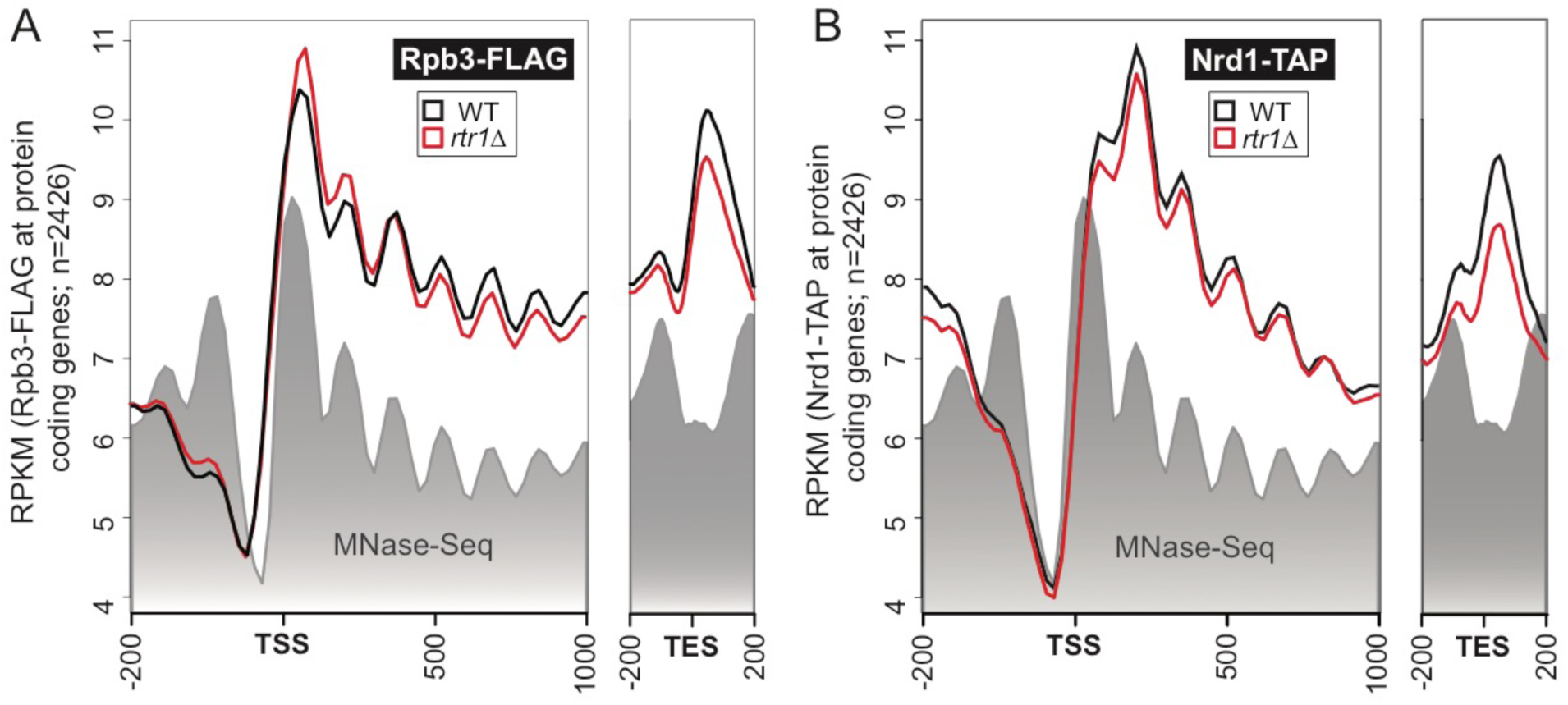
Deletion of *RTR1* decreases RNAPII occupancy through increased premature transcription termination at protein-coding genes. Average gene analysis of RNAPII (Rpb3) (A) or Nrd1 (B) levels at 2,426 protein-coding genes −200 nucleotides upstream and +1000 downstream of the TSS and −200/+200 upstream/downstream of the annotated TES. The data is shown as average RPKM values calculated using ngs.plot for each genotype designated by the colors in the legend.

The *IMD2* gene is regulated by an intergenic NNS-terminator which maintains basal levels of *IMD2* expression until low nucleotide levels stimulate *IMD2* expression (Figure 6A). In rich media conditions, such as those used for these experiments, an upstream *IMD2* CUT is produced and terminated by NNS-dependent termination. By ChIP-exo, both RNAPII and Nrd1 can be mapped to the upstream CUT region in WT cells with low levels of RNAPII observed in the *IMD2* coding region (Fig. 6B & C). However, both RNAPII and Nrd1 levels are significantly reduced in *rtr1Δ* cells similar to observations at snRNA genes (Fig. 6B & C). In addition to RNA-Seq analysis of WT and *rtr1Δ* cells we also performed RNA-Seq analysis of *rrp6Δ and rtr1Δ rrp6Δ* cells to determine the role of the Rrp6-containing exosome in the downregulation of ncRNA transcripts in *rtr1Δ* cells (Fig. S3A & B). The *IMD2* upstream CUT RNA is also decreased in *rtr1Δ* while it is increased in *rrp6Δ* (Fig. 6D). The *rtr1Δ rrp6Δ* cells show similar levels of the *IMD2* upstream CUT relative to WT suggesting that *rrp6Δ* is required to degrade the CUT following NNS-termination in *rtr1Δ* cells. The *IMD2* coding region is also altered by *rtr1Δ* perhaps due to the close proximity of Nrd1-Nab3 binding sites to the *IMD2* TSS (Fig. 6E). *IMD2* mRNA levels are ∼5-fold decreased in *rtr1Δ* cells but are not significantly impacted by *rrp6Δ* with *rtr1Δ rrp6Δ* cells also showing decreased levels relative to WT cells (Fig. 6E). Northern blot analysis of the *IMD2* coding region reflected similar ratios to RNA-Seq and confirmed that the terminator over-ride mutant in the Ser5-P CTD phosphatase Ssu72 (*ssu72-tov*) also showed increased expression of the full length *IMD2* transcript (Fig. 6F, [33]). These data suggest that while Ssu72 and Rrp6 are required for proper *IMD2* CUT termination and likely degradation, Rtr1 is a negative regulator of NNS termination with knockout of *RTR1* leading to decreased *IMD2* levels.

**Figure 6:**
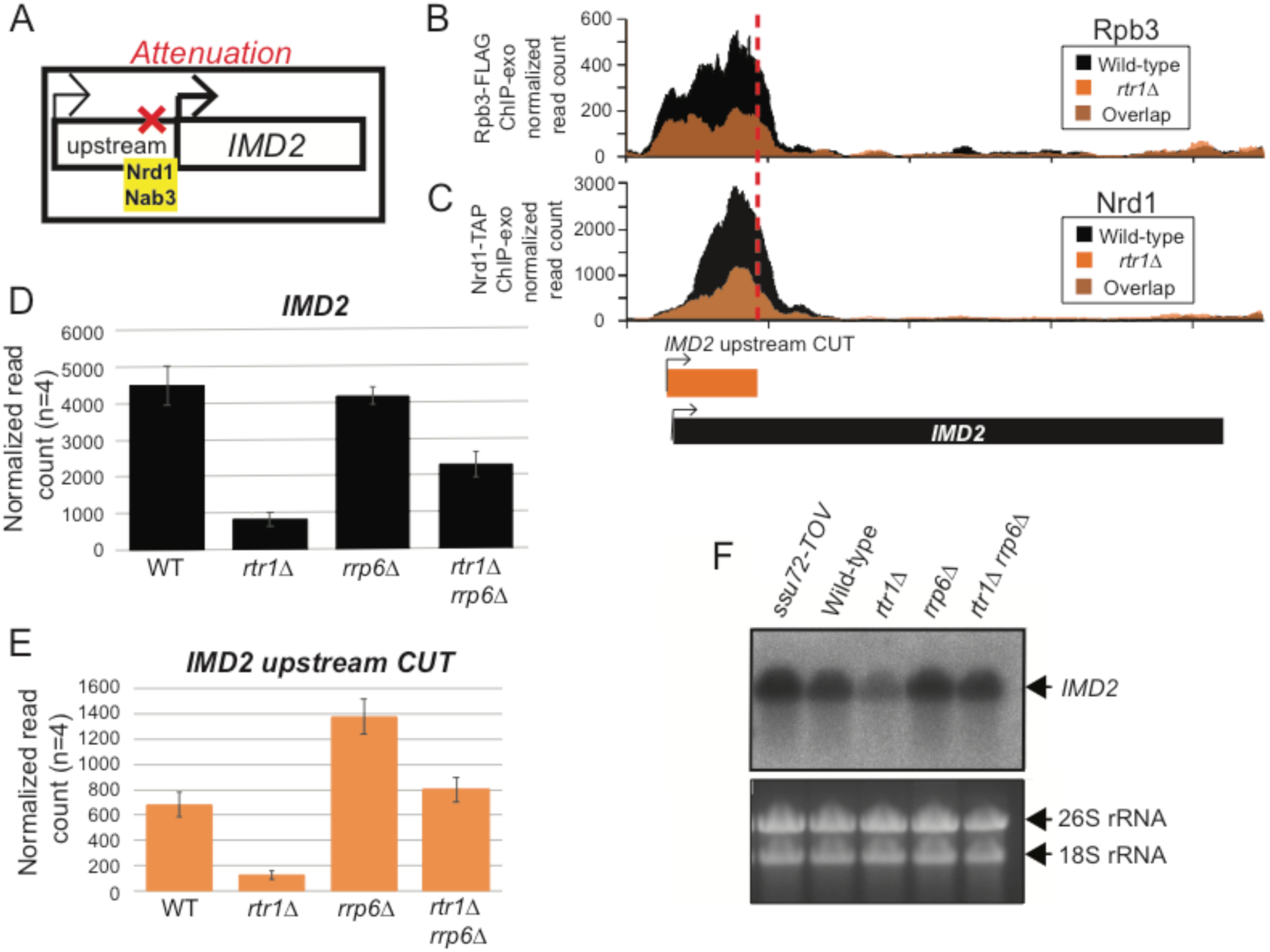
Rtr1 is required for basal *IMD2* expression. Occupancy of RNAPII (A) and Nrd1 (B) at the *IMD2* gene. Data derived from WT cells is in black, and those from the *rtr1Δ* strain are in orange. The location and direction of transcription for all analyzed annotations are diagrammed below the graphs, each to scale. C) Schematic representation of NNS-dependent attenuation of the *IMD2* gene. D) Expression of the *IMD2* upstream CUT by RNA-Seq (n=4). E) Expression of *IMD2* mRNA by RNA-Seq (n=4). F) Northern blot analysis of *IMD2* expression in various genotype cells as indicated.

### *Rtr1* promotes elongation of antisense noncoding transcripts

Since loss of the Rrp6-containing exosome in *rtr1Δ* leads to recovery of *IMD2* transcript levels, it is likely that Rtr1 is a positive regulator of noncoding RNA elongation in WT cells through attenuation of NNS-dependent termination. To explore this possibility, a comparison of global RNA-Seq analysis was performed through analysis of *rrp6Δ and rtr1Δ rrp6Δ* transcriptomes along with WT and *rtr1Δ* (Figure S4, Table S4). A focus on the antisense transcripts (ASTs) that were significantly downregulated in *rtr1Δ* relative to WT (n=104) revealed that the vast majority of the Rtr1-regulated ASTs depend on the Rrp6-containing exosome for their downregulation (Fig. 7). These data further suggest that the downregulation of the Rtr1-regulated ASTs is likely related to increased premature ncRNA termination through the NNS pathway which requires the Rrp6-containing exosome for RNA degradation [69]. Elongation of noncoding transcripts, including ASTs, has been shown to regulate protein coding gene expression through transcription interference [74]. Upregulation of noncoding transcript elongation, promoted by Rtr1 in WT cells, may therefore alter overall transcriptome profiles. It has been shown that transcription of the protein-coding gene *YKL151C* is regulated by an NNS-terminated antisense transcript, which is labelled as *YKL151C AS*. Elongation of the *YKL151C AS* leads to transcription interference of *YKL151C* expression (Fig. 8A). Northern blot analysis of *YKL151C / YKL151C AS* expression using single stranded RNA probes shows that *YKL151C* sense is up-regulated in *rtr1Δ* as a consequence of *YKL151C AS* transcript downregulation (Fig. 8B). The opposite effect can be observed in the *rrp6Δ,* and *rtr1Δ rrp6Δ* strains showing that decreased NNS-dependent termination leads to *YKL151C* sense elongation (Fig. 8). In *ssu72-tov* cells, *YKL151C* sense transcripts are decreased as well with a coordinate increase in *YKL151C* antisense although both changes occur to a lower extent than the changes seen in *rrp6Δ* (Fig. 8B). Furthermore, strand specific RNA-Seq analysis confirmed that *YKL151C* is upregulated in *rtr1Δ* while the *YKL151C AS* is significantly downregulated (Fig. 8C & D).

**Figure 7:**
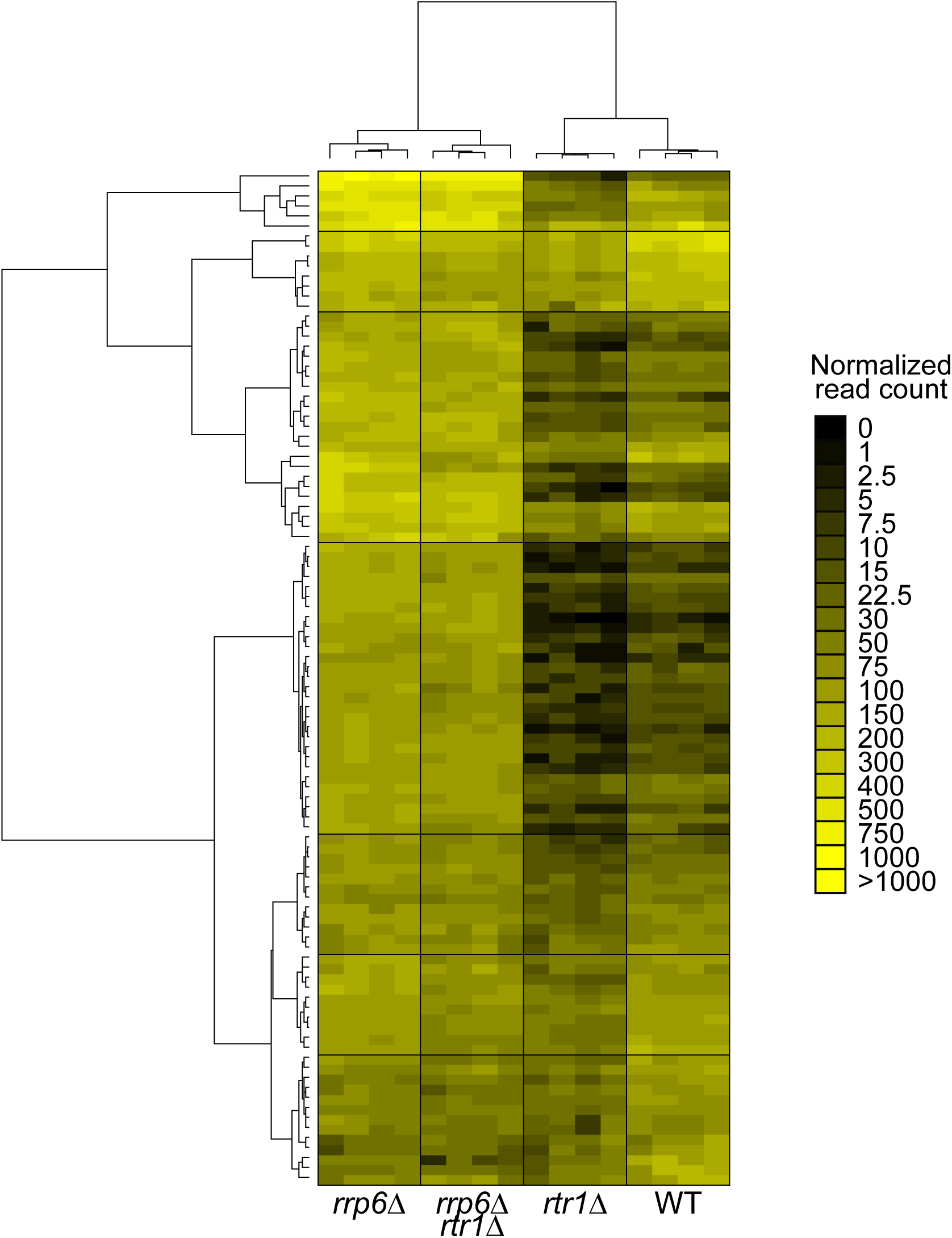
*RTR1/RRP6* double deletion strain phenocopies *rrp6Δ* cells. Heat map of transcripts differentially expressed in *rtr1Δ, rrp6Δ,* and *rtr1Δ rrp6Δ* compared to WT cells, according to the scale at the right. Unsupervised hierarchical clustering analysis was performed using normalized read count values for each biological replicate and genotype (n=16). Each genotype clustered together illustrating a high degree of reproducibility between biological replicates with WT values groups as Cluster IV, *rtr1Δ* values grouped as Cluster III, *rtr1Δ rrp6Δ* grouped as Cluster II, and *rrp6Δ* grouped as Cluster I.

**Figure 8:**
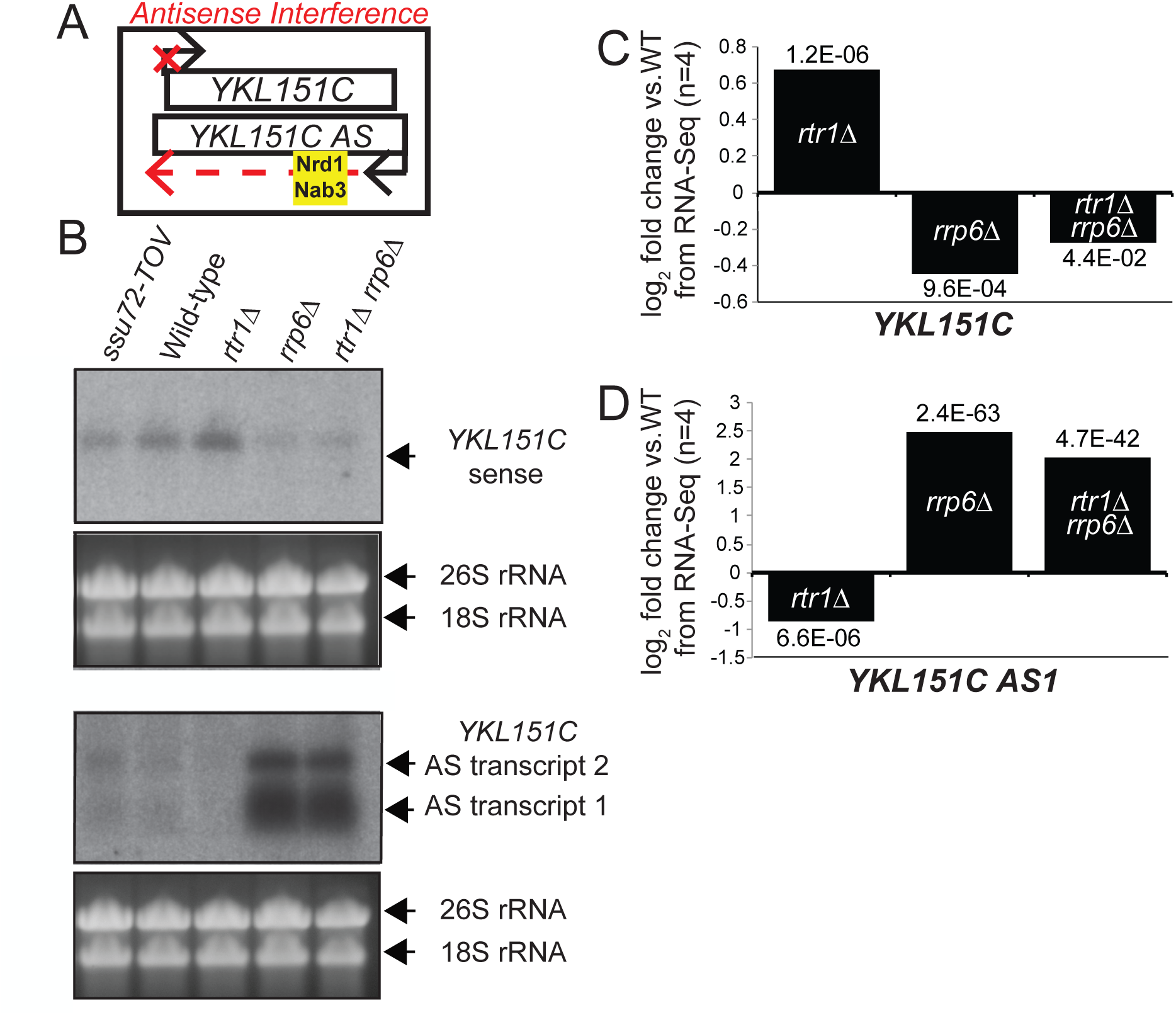
Analysis of YKL151C sense and antisense expression in WT and *rtr1Δ* cells. A) Schematic representation of *YKL151C* sense and antisense expression. B) Northern blot analysis of *YKL151C* sense and antisense transcripts using single-stranded RNA probes. C and D) Summary of RNA-Seq expression analysis of *YKL151C* sense and antisense transcript expression.

## Discussion

Our findings are the first to indicate that the Ser5 CTD phosphatase Rtr1 plays a role in premature termination of RNAPII transcripts through increasing the frequency of NNS-dependent termination. Our data indicate that Rtr1 normally limits the co-occupancy of Nrd1/RNAPII in WT cells, thereby acting as a key regulator of the NNS-dependent termination pathway and the balance between protein-coding and noncoding or cryptic transcription. In addition, these findings suggest that Rtr1 serves as a post-initiation intermediary facilitating the transition from early elongation to late elongation, likely promoting changes in cross-interactions between different transcription elongation and/or termination complexes. These findings, along with other recent reports, clearly establish Rtr1 as a general reader and eraser of the CTD code during RNAPII transcription elongation [48, 75-77].

DisCo network analysis was able to differentiate changes in protein complex interactions that occur between RNAPII, CFIa, CPF and NNS as a consequence of *RTR1* deletion. The decreased interaction between CFIa and RNAPII was evident through both prey-prey correlation analysis and SAINT probability analysis which also revealed that interactions between CFIa and CPF were detected less frequently in *rtr1Δ* cells. Additionally, interactions between Nrd1 and RNAPII were increased in cells lacking Rtr1 activity supporting our transcriptome-level findings that show that RNA produced from NNS-dependent genes can terminate earlier (Fig. 4) and more efficiently (Fig. 4-8). Overall, the *rtr1Δ* transcriptome has a clear decrease in gene expression relative to WT with a bias towards decreases in non-ORF transcripts (Fig. 2). These findings support models suggesting that the CTD is instrumental in the fate of termination choice and adds novel insights suggesting that Rtr1 is a major determinant that regulates alterations in transcription elongation-termination balance. Rtr1 and its human homolog RPAP2 have been implicated in the response to a variety of cellular stresses including changes in carbon source, heat, and ER stress inducers [78-80]. In light of the findings from this study, this suggest that Rtr1-dependent control of premature RNAPII termination could serve as a regulatory point to mitigate cellular stress.

This work also provides some new insights regarding the mechanisms of Nrd1-dependent RNAPII termination. As shown in Figure 3, RNAPII accumulates at well-defined Nrd1 binding sites such, in some cases, trailing RNAPII molecules pile up in some cases behind the Nrd1-bound RNAPII (Fig. 3C & D). This suggests that Nrd1 may cause RNAPII pausing as a consequence of RNA binding or perhaps coordinate RNA and RNAPII CTD binding. Furthermore, the affinity purification mass spectrometry studies suggest that Sen1 association with Nrd1-Nab3 occurs at a relatively low frequency. As discussed, SAINT probabilities of proteins that exist within a stable protein complex often exceed a value of 0.95 (Figure 1, [57, 75, 81]). These quantitative findings indicate that Sen1 is not a stable subunit of the Nrd1-Nab3 complex with SAINT probabilities suggesting that it is a transient interaction partner similar to the exosome and TRAMP complexes. A wealth of previous work has established the requirement for Sen1 in addition to Nrd1-Nab3 for RNAPII termination at sn/snoRNA genes *in vivo* (reviewed in [51]). Taken together, these data strongly suggest that Sen1 is recruited and/or diffuses transiently to terminate Nrd1-Nab3 paused RNAPII rather than functioning as an integral subunit of an NNS complex. *In vitro* studies have also shown that Sen1 is able to terminate RNAPII in the absence of Nrd1 and Nab3 further suggesting that Sen1 could potentially terminate multiple forms of paused RNAPII.

Through analysis of DisCo networks, RNA-Seq, and ChIP-exo, this work shows that Nrd1/RNAPII co-localization and protein-protein interaction is increased and NNS-dependent termination is enhanced in the absence of the atypical phosphatase Rtr1. This would suggest that Rtr1 attenuates the NNS pathway whereas the Ssu72 CTD phosphatase stimulates termination through NNS. This represents a novel role for Rtr1-dependent CTD dephosphorylation in the regulation of RNAPII termination choice in eukaryotes. Additionally, loss of *RTR1* is shown to have the opposite phenotype as the *RRP6* deletion at many NNS-target transcripts, and *rtr1Δ rrp6Δ* phenocopies the *rrp6Δ* strain. These findings support the hypothesis that Rtr1 helps to fine-tune NNS-dependent termination of transcription and subsequent termination-coupled RNA decay through the Rrp6-containing exosome. Overall, these findings suggest that the balance of premature termination vs. elongation of both coding and non-coding transcripts alters the steady state levels of different transcript classes allowing for both coarse and fine-tuned regulation of the transcriptome.

## Methods

### Yeast Strains

All yeast strains used are isogenic to BY4741. *RTR1* was knocked out of wild type and the *RRP6* deletion strain by homologous recombination with a kanamycin cassette to create the *RTR1* deletion and *RTR1/RRP6* double deletion strains. *RRP6* deletion strain is from the yeast knockout collection (Open Biosystems) [82]. The Rpb3-3xFLAG (referred to as Rpb3-FLAG) WT strain has been previously described [63]. The Nrd1-TAP strain is from the yeast TAP-tag collection (Open Biosystems). The Rpb3-FLAG and Nrd1-TAP *RTR1* deletion strains were made by amplification of the *RTR1* knockout cassette from the *RTR1* deletion strain and transformation into the wild-type Rpb3-FLAG and Nrd1-TAP strains respectively. All deletion strains were confirmed by PCR-based genotyping. To perform a single biological replicate for genomics or proteomics experiments, growths of all strains of interest were pre-cultured from a single colony obtained from a sequence-verified glycerol stock of the strain that had been plated on the appropriate selective medium and grown for 2 days. Liquid cultures of all genotypes for an individual biological experiment were grown up on the same day. Cells for subsequent biological replicates were grown on different days.

### Affinity purification of protein complexes

Cells were grown to OD600 ≅ 1.5 in YPD broth overnight and collected by centrifugation for 10 minutes at 4000 x g, then washed in H2O and resuspended in 25mL TAP lysis buffer per 2.5 grams of pellet (40mM Hepes-KOH, pH 7.5; 10% glycerol; 350mM NaCl; 0.1% Tween-20; fresh yeast protease inhibitors (Sigma; diluted to 1X)). The cells were slowly transferred to liquid nitrogen using a syringe. The frozen cells were pulverized with a mortar and pestle and lysed further in a Waring Blender with dry ice. The frozen lysate was transferred to a new container and allowed to thaw at room temperature. The resulting extract was treated with 100units DNase I and 10µL of 30mg/mL heparin for 10 minutes at room temperature and clarified by centrifugation as previously described [58]. Tandem Affinity Purification (TAP) was performed as previously described [58]. For FLAG tagged purifications, the lysate was incubated with anti-FLAG agarose resin (Sigma) at 4°C overnight. The resin and bound proteins were removed from the lysate by gravity flow through a 30mL Bio-Rad Econoprep column and washed on the column with 60 mL TAP lysis buffer. The resin was resuspended 300µL of 50mM Ammonium bicarbonate pH 8.0 and transfer to a microcentrifuge tube for on bead digestion with 5µL of Trypsin Gold (0.1µg/µL) overnight with shaking at 37°C. The supernatant containing the digested proteins was removed and treated with 20µL of 90% formic acid to inactivate the trypsin.

### MudPIT-LC/MS/MS and Proteomics Data Analysis

Each affinity purified sample was loaded onto a two-phase MudPIT column containing strong cation exchange resin (Phenomenex), which binds positively charged ions, and reverse phase C18 resin (Phenomenex), which will retain peptides based on their hydrophobicity [83]. The samples were eluted off the column by the MudPIT protocol of 10 steps of increasing salt concentrations (50-350mM ammonium acetate) followed by an organic gradient (20-80% acetonitrile). All chromatography solutions also contained 1% formic acid. Peptides were analyzed by a ThermoFisher LTQ Velos for MS/MS analysis. Raw spectrum data from the MS analysis were submitted for protein identification by Proteome Discoverer software (Thermo) version 2.1 using SEQUEST-HT as the database search algorithm. Database searches were performed against a FASTA database from the yeast Uniprot proteome. The FASTA database also included a number of common protein contaminants such as keratins and IgGs.

### Disruption – Compensation (DisCo) network analysis

DisCo analysis using protein-protein interaction tools to analyze protein-protein interaction dynamics as a consequence of genetic perturbation, in this case deletion of the CTD phosphatase *RTR1*. Statistical analysis of interactome (SAINT) was performed as previously described on at least four biological replicate purifications from each genotype [53, 54, 75]. In brief, PSMs for each copurfied protein were annotated per purification by bait protein, genotype (WT or ‘rtr1D’ for *rtr1Δ*), replicate in the list format used for analysis through crapome.org [84]. SAINTexpress was used for the probability score calculation [54]. The output file from SAINT analysis was used as the input for ProHits-viz which was employed for prey-prey correlation analysis with the following key options: Abundance column = Spec (i.e. PSM), Score column = Saint score, Abundance cutoff for prey correlation = 20, Add bait counts = yes [85].

### RNA Isolation

RNA was extracted using the hot acid phenol method described previously [63]. An Ambion DNase-turbo kit was used to degrade any contaminating DNA if the RNA was to be used for subsequent sequencing or PCR. The quality of the total RNA samples was determined with an Agilent Bioanalyzer before preparation of the sequencing libraries.

### Illumina HiSeq 4000 sequencing methods

Illumina TruSeq total RNA standard methods were used for yeast whole transcriptome sequencing. Total RNA was isolated and DNase treated (Ambion DNase). RNA was evaluated for quantity and quality for a minimum RIN score of 7 or higher using Agilent Bioanalyzer 2100. RNA samples were spiked with ERCC ExFold RNA spike-in mix (Life Technologies, 4456739) prior to library preparation. Samples were depleted of Ribosomal RNA using Ribo-Zero Magnetic Gold Kit (Illumina, MRZY1324). cDNA libraries were prepared using RNA fragmentation, cDNA synthesis, ligation of index adaptors, and amplification as specified in TruSeq sample preparation guide (Illumina, 15031048). Total RNA was sequenced with the Illumina HiSeq 4000. Ggplot was used for volcano plot visualization [86].

### ChIP-exo

Chromatin IP followed by exonuclease treatment was performed using the protocol described by Rhee and Pugh with the following specifics [66]. Rpb3-FLAG WT and *rtr1Δ* and Nrd1-TAP WT and *rtr1Δ* cells were grown to an OD_600_=0.8-1 prior to crosslinking with formaldehyde. Immunoprecipitation was performed with 50µL of anti-FLAG agarose or anti-TAP sepharose beads (Sigma). The volume of beads used for immunoprecipitation was optimized by affinity purification followed by mass spectrometry (data not shown). Subsequent sample processing steps including exonuclease treatment and sequencing library preparation were performed as previously described [66].

### MNase-Seq

Micrococcal nuclease (MNase) digest and sequencing was performed through adaptation of the protocol by Wal and Pugh [87]. Following optimization of the digestion conditions, 15U of MNase was added to a chromatin slurry and incubated with shaking at 37 °C for 20 minutes. The digestion was quenched by addition of 50 mM EDTA and 0.2% SDS. The digested DNA was cleaned up through phenol/chorloform extraction followed by ethanol precipitation with 20ug of glycogen (Sigma) as a carrier.

### SOLiD5500xl sequencing methods

ChIP-exo and MNase-Seq library construction, EZBead preparation, and Next-Gen sequencing were completed using standard methods based on the Life Technologies SOLiD5500xl system as previously described [63]. The resulting 75 nt solid reads were mapped to *Saccharomyces cerevisiae* sacCer3 reference genome using in-house mapping pipelines that utilizes bfast-0.7.0a [88]. First, rRNA, tRNAs, and poor-quality reads were discarded, and remaining reads were mapped to reference genome sacCer3 and a splice-junction library, respectively. The genomic and splice-junction library mapping data were merged at the end. In a second pipeline, the rRNA/tRNA reads were kept and mapped to the reference genome only due to the minimal number of introns in the yeast genome. Read counts per nucleotide were calculated using bamutils from NGSUtils [89]. Differential gene expression was analyzed using edgeR, which has been shown to work well with low replicate numbers [60, 90]. Four biological replicates were used for each genotype in the RNA-Seq analysis. All raw and processed files from the RNA sequencing and ChIP-exo experiments performed for this study have been deposited to Gene Expression Omnibus [GEO] under the accession number GSE87657.

### Genomics data analysis

Following data alignment, noncoding transcripts were manually inspected individually using the Integrative Genomics Viewer [91]. To identify ASTs with significant changes in differential expression, the strand was reversed for all sense annotations for the coding region of each ORF-Ts and the text “AS_” was added in front of the ORF-T name. The annotations for the 5’ and 3’ UTR were not included. These annotations were then used for edgeR analysis and the annotations for ASTs that showed significant changes in *rrp6Δ* were used for subsequent differential expression analysis to generate the final dataset in Supplementary Table S1. The average gene analysis plots for different RNAPII gene classes were generated using data from two biological replicate experiments per genotype per plot with the program ngs.plot using data from bam files and further edited in Adobe Illustrator [73]. The plots include the standard error of the mean for the total number of genes (defined in the text and figure legends) used for average gene analysis calculated by ngs.plot.

### Northern Blot Analysis

Northern blot analysis was performed as previously described [63]. Thirty micrograms of total RNA were loaded per lane on a 1% agarose gel and separated by electrophoresis at 120 volts for 1 hour at 4°C. The RNA was transferred to Bio-Rad Zeta-Probe® blotting membranes by capillary overnight. Transfer efficiency was determined by Methylene Blue staining. Strand specific RNA probes were expressed from a linearized pET-DEST42 (Invitrogen) containing the region of interest in the sense or antisense orientation by T7 transcription (MAXIscript) using ^32^P labeled UTP. The radiolabeled probe was purified and then hybridized to the RNA blot at 68°C overnight. The membranes were then washed with 1xSSC/.1%SDS twice at room temperature and twice with .1xSSC/.1%SDS for 15 minutes at 68°C. Blots were exposed to a phosphorscreen followed by scanning using a phosphorimager (GE Healthcare).

## ACKNOWLEDGMENTS

The authors would like to thank the members of the Mosley lab and Jerry Workman for critical reading and discussion of the data in this manuscript. We would also like to thank Joseph Bidwell and Ronald Wek for helpful discussions of this work. The *ssu72 TOV* yeast strain was kindly provided by the Reines lab. We would like to sincerely thank Howard Edenberg, Xiaoling Xuei, and the other members of the IU School of Medicine Center for Medical Genomics for their advice and efforts in the sequencing studies included in this work. Funding for the majority of this work was provided by National Institutes of Health grant R01GM099714 to A.L.M. including supplement funding to support J.F.V. The development of the DisCo network approach was supported by the National Science Foundation award 1515748 to A.L.M. The initial phases of this study were also supported by the Showalter Research Trust.

## Supporting Information Legends

**Table S1: Peptide – spectrum match (PSM) table for the termination factor purification dataset.** Protein names are given in the ProtID column according to their Uniprot ID. Each subsequent column represents an individual biological sample with the bait name and genotype defined in the column header. BY4741 is the isogenic parental yeast strain for all mutants used in this study and was used for mock control purifications.

**Table S2: Statistical Analysis of INTeractome (SAINT) dataset for the termination factor purifications.** Standard fold-change values are calculated using the average PSM values from the controls whereas stringent fold-change values were calculated using the maximum PSM values across all controls performed by SAINT express [1]. The bait name and genotype are given in the column header. The iREF values equal to 1 indicates that the bait and the protein have previously been described as interacting proteins in previous work.

**Table S3: *RTR1* knockout (KO) cell transcriptome data relative to WT.** Table with fold-change, p-value, and false discovery rate (FDR) calculated by edgeR. Sum, average, and individual biological replicate (Rep n) normalized read counts for WT and *RTR1* deletion data [2].

**Table S4: RRP6 knockout (KO) and RTR1 RRP6 knockout cell transcriptome data relative to WT.** Table with fold-change, p-value, and false discovery rate (FDR) calculated by edgeR. Sum, average, and individual biological replicate (Rep n) normalized read counts for WT and *RRP6* and *RTR1 RRP6* knockout data.

**Figure S1: STRING network analysis of termination factor complex data using a fold-change cutoff of 5 or more** [3]. Networks are included for Pcf11, Nrd1, and Ssu72 purifications from WT cells (BY4741). Figure legends are included for each network with a selection of enriched set of proteins defined using pathway analysis.

**Figure S2:** Enrichment of Nrd1 occupancy relative to total RNAPII at protein coding genes. Genes are sorted by increasing gene length when the annotated transcription end site TES defined by a dashed line at the 3’-end [4]. All genes are aligned at the 5’- end by the annotated transcription start site (TSS). Nrd1 levels are clearly depleted relative to RNAPII at the TSS followed by enriched levels of Nrd1 just downstream of the TSS.

**Figure S3: Average ChIP-exo occupancy profiles for RNAPII (Rpb3) and Nrd1 from WT and RTR1 knockout cells.** The legend defines the line color for each sample as indicated on the left. MNase-Seq based histone occupancy is also shown as gray shaded profiles [4].

**Figure S4: Volcano plots representing significant changes in the *RRP6* and *RRP6 RTR1* knockout transcriptomes relative to WT.** Density plots are included to illustrate the number of points in each area as indicated. The number of decreased and increased transcripts based on a fold-change cutoff of 1.5-fold and an FDR of at least 0.05 are shown at the top of each panel for *rrp6Δ* (A) and *rrp6Δ rtr1Δ* (B) [5].

## REFERENCES

1. Hazelbaker DZ, Marquardt S, Wlotzka W, Buratowski S. Kinetic competition between RNA Polymerase II and Sen1-dependent transcription termination. Mol Cell. 2013;49(1):55–66. Epub 2012/11/28. doi: 10.1016/j.molcel.2012.10.014. PubMed PMID: 23177741; PubMed Central PMCID: PMC3545030.

2. Kamieniarz-Gdula K, Proudfoot NJ. Transcriptional Control by Premature Termination: A Forgotten Mechanism. Trends Genet. 2019. Epub 2019/06/20. doi: 10.1016/j.tig.2019.05.005. PubMed PMID: 31213387.

3. Kamieniarz-Gdula K, Gdula MR, Panser K, Nojima T, Monks J, Wisniewski JR, et al. Selective Roles of Vertebrate PCF11 in Premature and Full-Length Transcript Termination. Mol Cell. 2019;74(1):158–72 e9. Epub 2019/03/02. doi: 10.1016/j.molcel.2019.01.027. PubMed PMID: 30819644; PubMed Central PMCID: PMCPMC6458999.

4. Schlackow M, Nojima T, Gomes T, Dhir A, Carmo-Fonseca M, Proudfoot NJ. Distinctive Patterns of Transcription and RNA Processing for Human lincRNAs. Mol Cell. 2017;65(1):25–38. Epub 2016/12/27. doi: 10.1016/j.molcel.2016.11.029. PubMed PMID: 28017589; PubMed Central PMCID: PMCPMC5222723.

5. Dhir A, Dhir S, Proudfoot NJ, Jopling CL. Microprocessor mediates transcriptional termination of long noncoding RNA transcripts hosting microRNAs. Nat Struct Mol Biol. 2015;22(4):319–27. Epub 2015/03/03. doi: 10.1038/nsmb.2982. PubMed PMID: 25730776; PubMed Central PMCID: PMCPMC4492989.

6. Wagschal A, Rousset E, Basavarajaiah P, Contreras X, Harwig A, Laurent-Chabalier S, et al. Microprocessor, Setx, Xrn2, and Rrp6 co-operate to induce premature termination of transcription by RNAPII. Cell. 2012;150(6):1147–57. Epub 2012/09/18. doi: 10.1016/j.cell.2012.08.004. PubMed PMID: 22980978; PubMed Central PMCID: PMCPMC3595997.

7. Fusby B, Kim S, Erickson B, Kim H, Peterson ML, Bentley DL. Coordination of RNA Polymerase II Pausing and 3’ End Processing Factor Recruitment with Alternative Polyadenylation. Molecular and cellular biology. 2016;36(2):295–303. Epub 2015/11/04. doi: 10.1128/MCB.00898-15. PubMed PMID: 26527620; PubMed Central PMCID: PMCPMC4719304.

8. Brody Y, Neufeld N, Bieberstein N, Causse SZ, Bohnlein EM, Neugebauer KM, et al. The in vivo kinetics of RNA polymerase II elongation during co-transcriptional splicing. PLoS Biol. 2011;9(1):e1000573. Epub 2011/01/26. doi: 10.1371/journal.pbio.1000573. PubMed PMID: 21264352; PubMed Central PMCID: PMCPMC3019111.

9. Liu X, Freitas J, Zheng D, Oliveira MS, Hoque M, Martins T, et al. Transcription elongation rate has a tissue-specific impact on alternative cleavage and polyadenylation in Drosophila melanogaster. Rna. 2017;23(12):1807–16. Epub 2017/08/31. doi: 10.1261/rna.062661.117. PubMed PMID: 28851752; PubMed Central PMCID: PMCPMC5689002.

10. Corden JL. Tails of RNA polymerase II. Trends Biochem Sci. 1990;15(10):383–7. Epub 1990/10/01. PubMed PMID: 2251729.

11. Buratowski S. Progression through the RNA polymerase II CTD cycle. Mol Cell. 2009;36(4):541–6. Epub 2009/11/28. doi: 10.1016/j.molcel.2009.10.019. PubMed PMID: 19941815; PubMed Central PMCID: PMC3232742.

12. Mosley AL, Pattenden SG, Carey M, Venkatesh S, Gilmore JM, Florens L, et al. Rtr1 is a CTD phosphatase that regulates RNA polymerase II during the transition from serine 5 to serine 2 phosphorylation. Mol Cell. 2009;34(2):168–78. Epub 2009/04/28. doi: 10.1016/j.molcel.2009.02.025. PubMed PMID: 19394294; PubMed Central PMCID: PMCPMC2996052.

13. Zhang DW, Mosley AL, Ramisetty SR, Rodriguez-Molina JB, Washburn MP, Ansari AZ. Ssu72 phosphatase-dependent erasure of phospho-Ser7 marks on the RNA polymerase II C-terminal domain is essential for viability and transcription termination. J Biol Chem. 2012;287(11):8541–51. Epub 2012/01/12. doi: 10.1074/jbc.M111.335687. PubMed PMID: 22235117; PubMed Central PMCID: PMC3318730.

14. Jeronimo C, Collin P, Robert F. The RNA Polymerase II CTD: The Increasing Complexity of a Low-Complexity Protein Domain. J Mol Biol. 2016;428(12):2607–22. Epub 2016/02/16. doi: 10.1016/j.jmb.2016.02.006. PubMed PMID: 26876604.

15. Bataille AR, Jeronimo C, Jacques PE, Laramee L, Fortin ME, Forest A, et al. A universal RNA polymerase II CTD cycle is orchestrated by complex interplays between kinase, phosphatase, and isomerase enzymes along genes. Mol Cell. 2012;45(2):158–70. Epub 2012/01/31. doi: 10.1016/j.molcel.2011.11.024. PubMed PMID: 22284676.

16. Schreieck A, Easter AD, Etzold S, Wiederhold K, Lidschreiber M, Cramer P, et al. RNA polymerase II termination involves C-terminal-domain tyrosine dephosphorylation by CPF subunit Glc7. Nat Struct Mol Biol. 2014;21(2):175–9. Epub 2014/01/15. doi: 10.1038/nsmb.2753. PubMed PMID: 24413056; PubMed Central PMCID: PMCPMC3917824.

17. Nedea E, He X, Kim M, Pootoolal J, Zhong G, Canadien V, et al. Organization and function of APT, a subcomplex of the yeast cleavage and polyadenylation factor involved in the formation of mRNA and small nucleolar RNA 3’-ends. J Biol Chem. 2003;278(35):33000–10. Epub 2003/06/24. doi: 10.1074/jbc.M304454200. PubMed PMID: 12819204.

18. Kong SE, Kobor MS, Krogan NJ, Somesh BP, Sogaard TM, Greenblatt JF, et al. Interaction of Fcp1 phosphatase with elongating RNA polymerase II holoenzyme, enzymatic mechanism of action, and genetic interaction with elongator. J Biol Chem. 2005;280(6):4299–306. Epub 2004/11/26. doi: 10.1074/jbc.M411071200. PubMed PMID: 15563457.

19. Kobor MS, Archambault J, Lester W, Holstege FC, Gileadi O, Jansma DB, et al. An unusual eukaryotic protein phosphatase required for transcription by RNA polymerase II and CTD dephosphorylation in S. cerevisiae. Mol Cell. 1999;4(1):55–62. Epub 1999/08/13. PubMed PMID: 10445027.

20. Schwer B, Ghosh A, Sanchez AM, Lima CD, Shuman S. Genetic and structural analysis of the essential fission yeast RNA polymerase II CTD phosphatase Fcp1. Rna. 2015;21(6):1135–46. Epub 2015/04/18. doi: 10.1261/rna.050286.115. PubMed PMID: 25883047; PubMed Central PMCID: PMCPMC4436666.

21. Ghosh A, Shuman S, Lima CD. The structure of Fcp1, an essential RNA polymerase II CTD phosphatase. Mol Cell. 2008;32(4):478–90. Epub 2008/11/26. doi: 10.1016/j.molcel.2008.09.021. PubMed PMID: 19026779; PubMed Central PMCID: PMCPMC2645342.

22. Hunter GO, Fox MJ, Smith-Kinnaman WR, Gogol M, Fleharty B, Mosley AL. Phosphatase Rtr1 Regulates Global Levels of Serine 5 RNA Polymerase II C-Terminal Domain Phosphorylation and Cotranscriptional Histone Methylation. Molecular and cellular biology. 2016;36(17):2236–45. Epub 2016/06/02. doi: 10.1128/MCB.00870-15. PubMed PMID: 27247267; PubMed Central PMCID: PMCPMC4985930.

23. Hsu PL, Yang F, Smith-Kinnaman W, Yang W, Song JE, Mosley AL, et al. Rtr1 is a dual specificity phosphatase that dephosphorylates Tyr1 and Ser5 on the RNA polymerase II CTD. J Mol Biol. 2014;426(16):2970–81. Epub 2014/06/22. doi: 10.1016/j.jmb.2014.06.010. PubMed PMID: 24951832; PubMed Central PMCID: PMCPMC4119023.

24. Krishnamurthy S, He X, Reyes-Reyes M, Moore C, Hampsey M. Ssu72 Is an RNA polymerase II CTD phosphatase. Mol Cell. 2004;14(3):387–94. Epub 2004/05/06. PubMed PMID: 15125841.

25. Meinhart A, Silberzahn T, Cramer P. The mRNA transcription/processing factor Ssu72 is a potential tyrosine phosphatase. J Biol Chem. 2003;278(18):15917–21. Epub 2003/02/28. doi: 10.1074/jbc.M301643200. PubMed PMID: 12606538.

26. Lunde BM, Reichow SL, Kim M, Suh H, Leeper TC, Yang F, et al. Cooperative interaction of transcription termination factors with the RNA polymerase II C-terminal domain. Nat Struct Mol Biol. 2010;17(10):1195–201. Epub 2010/09/08. doi: 10.1038/nsmb.1893. PubMed PMID: 20818393; PubMed Central PMCID: PMC2950884.

27. Vasiljeva L, Kim M, Mutschler H, Buratowski S, Meinhart A. The Nrd1-Nab3-Sen1 termination complex interacts with the Ser5-phosphorylated RNA polymerase II C-terminal domain. Nat Struct Mol Biol. 2008;15(8):795–804. Epub 2008/07/29. doi: 10.1038/nsmb.1468. PubMed PMID: 18660819; PubMed Central PMCID: PMCPMC2597375.

28. Kim M, Vasiljeva L, Rando OJ, Zhelkovsky A, Moore C, Buratowski S. Distinct pathways for snoRNA and mRNA termination. Mol Cell. 2006;24(5):723–34. Epub 2006/12/13. doi: 10.1016/j.molcel.2006.11.011. PubMed PMID: 17157255.

29. Vasiljeva L, Buratowski S. Nrd1 interacts with the nuclear exosome for 3’ processing of RNA polymerase II transcripts. Mol Cell. 2006;21(2):239–48. Epub 2006/01/24. doi: 10.1016/j.molcel.2005.11.028. PubMed PMID: 16427013.

30. Creamer TJ, Darby MM, Jamonnak N, Schaughency P, Hao H, Wheelan SJ, et al. Transcriptome-wide binding sites for components of the Saccharomyces cerevisiae non-poly(A) termination pathway: Nrd1, Nab3, and Sen1. PLoS Genet. 2011;7(10):e1002329. Epub 2011/10/27. doi: 10.1371/journal.pgen.1002329. PubMed PMID: 22028667; PubMed Central PMCID: PMC3197677.

31. Jamonnak N, Creamer TJ, Darby MM, Schaughency P, Wheelan SJ, Corden JL. Yeast Nrd1, Nab3, and Sen1 transcriptome-wide binding maps suggest multiple roles in post-transcriptional RNA processing. Rna. 2011;17(11):2011–25. Epub 2011/09/29. doi: 10.1261/rna.2840711. PubMed PMID: 21954178; PubMed Central PMCID: PMCPMC3198594.

32. Arigo JT, Carroll KL, Ames JM, Corden JL. Regulation of yeast NRD1 expression by premature transcription termination. Mol Cell. 2006;21(5):641–51. Epub 2006/03/02. doi: 10.1016/j.molcel.2006.02.005. PubMed PMID: 16507362.

33. Loya TJ, O’Rourke TW, Reines D. A genetic screen for terminator function in yeast identifies a role for a new functional domain in termination factor Nab3. Nucleic Acids Res. 2012;40(15):7476–91. Epub 2012/05/09. doi: 10.1093/nar/gks377. PubMed PMID: 22564898; PubMed Central PMCID: PMCPMC3424548.

34. Jenks MH, O’Rourke TW, Reines D. Properties of an intergenic terminator and start site switch that regulate IMD2 transcription in yeast. Molecular and cellular biology. 2008;28(12):3883–93. Epub 2008/04/23. doi: 10.1128/MCB.00380-08. PubMed PMID: 18426909; PubMed Central PMCID: PMCPMC2423123.

35. Kopcewicz KA, O’Rourke TW, Reines D. Metabolic regulation of IMD2 transcription and an unusual DNA element that generates short transcripts. Molecular and cellular biology. 2007;27(8):2821–9. Epub 2007/02/14. doi: 10.1128/MCB.02159-06. PubMed PMID: 17296737; PubMed Central PMCID: PMCPMC1899919.

36. Grzechnik P, Gdula MR, Proudfoot NJ. Pcf11 orchestrates transcription termination pathways in yeast. Genes Dev. 2015;29(8):849–61. Epub 2015/04/17. doi: 10.1101/gad.251470.114. PubMed PMID: 25877920; PubMed Central PMCID: PMCPMC4403260.

37. Mayer A, Heidemann M, Lidschreiber M, Schreieck A, Sun M, Hintermair C, et al. CTD tyrosine phosphorylation impairs termination factor recruitment to RNA polymerase II. Science. 2012;336(6089):1723–5. Epub 2012/06/30. doi: 10.1126/science.1219651. PubMed PMID: 22745433.

38. Kim M, Krogan NJ, Vasiljeva L, Rando OJ, Nedea E, Greenblatt JF, et al. The yeast Rat1 exonuclease promotes transcription termination by RNA polymerase II. Nature. 2004;432(7016):517–22. Epub 2004/11/27. doi: 10.1038/nature03041. PubMed PMID: 15565157.

39. Fong N, Brannan K, Erickson B, Kim H, Cortazar MA, Sheridan RM, et al. Effects of Transcription Elongation Rate and Xrn2 Exonuclease Activity on RNA Polymerase II Termination Suggest Widespread Kinetic Competition. Mol Cell. 2015;60(2):256–67. Epub 2015/10/17. doi: 10.1016/j.molcel.2015.09.026. PubMed PMID: 26474067; PubMed Central PMCID: PMCPMC4654110.

40. Pearson EL, Moore CL. Dismantling promoter-driven RNA polymerase II transcription complexes in vitro by the termination factor Rat1. J Biol Chem. 2013;288(27):19750–9. Epub 2013/05/22. doi: 10.1074/jbc.M112.434985. PubMed PMID: 23689372; PubMed Central PMCID: PMCPMC3707679.

41. Baejen C, Andreani J, Torkler P, Battaglia S, Schwalb B, Lidschreiber M, et al. Genome-wide Analysis of RNA Polymerase II Termination at Protein-Coding Genes. Mol Cell. 2017;66(1):38–49 e6. Epub 2017/03/21. doi: 10.1016/j.molcel.2017.02.009. PubMed PMID: 28318822.

42. Park J, Kang M, Kim M. Unraveling the mechanistic features of RNA polymerase II termination by the 5’-3’ exoribonuclease Rat1. Nucleic Acids Res. 2015;43(5):2625–37. Epub 2015/02/28. doi: 10.1093/nar/gkv133. PubMed PMID: 25722373; PubMed Central PMCID: PMCPMC4357727.

43. Proudfoot NJ. Transcriptional termination in mammals: Stopping the RNA polymerase II juggernaut. Science. 2016;352(6291):aad9926. Epub 2016/06/11. doi: 10.1126/science.aad9926. PubMed PMID: 27284201; PubMed Central PMCID: PMCPMC5144996.

44. Eaton JD, West S. An end in sight? Xrn2 and transcriptional termination by RNA polymerase II. Transcription. 2018;9(5):321–6. Epub 2018/07/24. doi: 10.1080/21541264.2018.1498708. PubMed PMID: 30035655; PubMed Central PMCID: PMCPMC6150625.

45. Fox MJ, Gao H, Smith-Kinnaman WR, Liu Y, Mosley AL. The exosome component Rrp6 is required for RNA polymerase II termination at specific targets of the Nrd1-Nab3 pathway. PLoS Genet. 2015;11(2):e1004999. Epub 2015/02/14. doi: 10.1371/journal.pgen.1004999. PubMed PMID: 25680078; PubMed Central PMCID: PMCPMC4378619.

46. Lemay JF, Bachand F. Fail-safe transcription termination: Because one is never enough. RNA Biol. 2015;12(9):927–32. Epub 2015/08/15. doi: 10.1080/15476286.2015.1073433. PubMed PMID: 26273910; PubMed Central PMCID: PMCPMC4615224.

47. Lemay JF, Larochelle M, Marguerat S, Atkinson S, Bahler J, Bachand F. The RNA exosome promotes transcription termination of backtracked RNA polymerase II. Nat Struct Mol Biol. 2014;21(10):919–26. Epub 2014/09/23. doi: 10.1038/nsmb.2893. PubMed PMID: 25240800.

48. Hunter GO, Fox MJ, Smith-Kinnaman WR, Gogol M, Fleharty B, Mosley AL. The phosphatase Rtr1 regulates global levels of serine 5 RNA Polymerase II C-terminal domain phosphorylation and cotranscriptional histone methylation. Molecular and cellular biology. 2016. doi: 10.1128/MCB.00870-15. PubMed PMID: 27247267.

49. Gudipati RK, Villa T, Boulay J, Libri D. Phosphorylation of the RNA polymerase II C-terminal domain dictates transcription termination choice. Nat Struct Mol Biol. 2008;15(8):786–94. Epub 2008/07/29. doi: 10.1038/nsmb.1460. PubMed PMID: 18660821.

50. Vasiljeva L, Kim M, Mutschler H, Buratowski S, Meinhart A. The Nrd1-Nab3-Sen1 termination complex interacts with the Ser5-phosphorylated RNA polymerase II C-terminal domain. Nature structural & molecular biology. 2008;15(8):795–804. Epub 2008/07/29. doi: 10.1038/nsmb.1468. PubMed PMID: 18660819; PubMed Central PMCID: PMC2597375.

51. Arndt KM, Reines D. Termination of Transcription of Short Noncoding RNAs by RNA Polymerase II. Annual review of biochemistry. 2015;84:381–404. doi: 10.1146/annurev-biochem-060614-034457. PubMed PMID: 25747400.

52. Porrua O, Libri D. Transcription termination and the control of the transcriptome: why, where and how to stop. Nature reviews Molecular cell biology. 2015;16(3):190–202. doi: 10.1038/nrm3943. PubMed PMID: 25650800.

53. Choi H, Larsen B, Lin ZY, Breitkreutz A, Mellacheruvu D, Fermin D, et al. SAINT: probabilistic scoring of affinity purification-mass spectrometry data. Nat Methods. 2011;8(1):70–3. Epub 2010/12/07. doi: 10.1038/nmeth.1541 nmeth.1541 [pii]. PubMed PMID: 21131968; PubMed Central PMCID: PMC3064265.

54. Teo G, Liu G, Zhang J, Nesvizhskii AI, Gingras AC, Choi H. SAINTexpress: improvements and additional features in Significance Analysis of INTeractome software. J Proteomics. 2014;100:37–43. Epub 2014/02/12. doi: 10.1016/j.jprot.2013.10.023S1874-3919(13)00538-1 [pii]. PubMed PMID: 24513533; PubMed Central PMCID: PMC4102138.

55. Szklarczyk D, Gable AL, Lyon D, Junge A, Wyder S, Huerta-Cepas J, et al. STRING v11: protein-protein association networks with increased coverage, supporting functional discovery in genome-wide experimental datasets. Nucleic Acids Res. 2019;47(D1):D607–D13. Epub 2018/11/27. doi: 10.1093/nar/gky1131. PubMed PMID: 30476243; PubMed Central PMCID: PMCPMC6323986.

56. Casanal A, Kumar A, Hill CH, Easter AD, Emsley P, Degliesposti G, et al. Architecture of eukaryotic mRNA 3’-end processing machinery. Science. 2017;358(6366):1056–9. Epub 2017/10/28. doi: 10.1126/science.aao6535. PubMed PMID: 29074584; PubMed Central PMCID: PMCPMC5788269.

57. Bedard LG, Dronamraju R, Kerschner JL, Hunter GO, Axley ED, Boyd AK, et al. Quantitative Analysis of Dynamic Protein Interactions during Transcription Reveals a Role for Casein Kinase II in Polymerase-associated Factor (PAF) Complex Phosphorylation and Regulation of Histone H2B Monoubiquitylation. J Biol Chem. 2016;291(26):13410–20. Epub 2016/05/05. doi: 10.1074/jbc.M116.727735. PubMed PMID: 27143358; PubMed Central PMCID: PMCPMC4919428.

58. Mosley AL, Sardiu ME, Pattenden SG, Workman JL, Florens L, Washburn MP. Highly reproducible label free quantitative proteomic analysis of RNA polymerase complexes. Mol Cell Proteomics. 2011;10(2):M110 000687. Epub 2010/11/05. doi: 10.1074/mcp.M110.000687 M110.000687 [pii]. PubMed PMID: 21048197; PubMed Central PMCID: PMC3033667.

59. Jiang L, Schlesinger F, Davis CA, Zhang Y, Li R, Salit M, et al. Synthetic spike-in standards for RNA-seq experiments. Genome Res. 2011;21(9):1543–51. Epub 2011/08/06. doi: 10.1101/gr.121095.111. PubMed PMID: 21816910; PubMed Central PMCID: PMCPMC3166838.

60. Robinson MD, McCarthy DJ, Smyth GK. edgeR: a Bioconductor package for differential expression analysis of digital gene expression data. Bioinformatics. 2010;26(1):139–40. Epub 2009/11/17. doi: 10.1093/bioinformatics/btp616. PubMed PMID: 19910308; PubMed Central PMCID: PMCPMC2796818.

61. Neil H, Malabat C, d’Aubenton-Carafa Y, Xu Z, Steinmetz LM, Jacquier A. Widespread bidirectional promoters are the major source of cryptic transcripts in yeast. Nature. 2009;457(7232):1038–42. Epub 2009/01/27. doi: 10.1038/nature07747. PubMed PMID: 19169244.

62. Schulz D, Schwalb B, Kiesel A, Baejen C, Torkler P, Gagneur J, et al. Transcriptome surveillance by selective termination of noncoding RNA synthesis. Cell. 2013;155(5):1075–87. Epub 2013/11/12. doi: 10.1016/j.cell.2013.10.024. PubMed PMID: 24210918.

63. Fox MJ, Gao H, Smith-Kinnaman WR, Liu Y, Mosley AL. The exosome component Rrp6 is required for RNA polymerase II termination at specific targets of the Nrd1-Nab3 pathway. PLoS Genet. 2015;10(2):e1004999. Epub 2015/02/14. doi: 10.1371/journal.pgen.1004999. PubMed PMID: 25680078.

64. Merran J, Corden JL. Yeast RNA-Binding Protein Nab3 Regulates Genes Involved in Nitrogen Metabolism. Molecular and cellular biology. 2017;37(18). Epub 2017/07/05. doi: 10.1128/MCB.00154-17. PubMed PMID: 28674185; PubMed Central PMCID: PMCPMC5574042.

65. Kuehner JN, Brow DA. Regulation of a eukaryotic gene by GTP-dependent start site selection and transcription attenuation. Mol Cell. 2008;31(2):201–11. Epub 2008/07/29. doi: 10.1016/j.molcel.2008.05.018. PubMed PMID: 18657503.

66. Rhee HS, Pugh BF. ChIP-exo method for identifying genomic location of DNA-binding proteins with near-single-nucleotide accuracy. Curr Protoc Mol Biol. 2012;Chapter 21:Unit 21 4. Epub 2012/10/03. doi: 10.1002/0471142727.mb2124s100. PubMed PMID: 23026909; PubMed Central PMCID: PMC3813302.

67. Thiebaut M, Colin J, Neil H, Jacquier A, Seraphin B, Lacroute F, et al. Futile cycle of transcription initiation and termination modulates the response to nucleotide shortage in S. cerevisiae. Mol Cell. 2008;31(5):671–82. Epub 2008/09/09. doi: 10.1016/j.molcel.2008.08.010. PubMed PMID: 18775327.

68. Vasiljeva L, Kim M, Terzi N, Soares LM, Buratowski S. Transcription termination and RNA degradation contribute to silencing of RNA polymerase II transcription within heterochromatin. Mol Cell. 2008;29(3):313–23. Epub 2008/02/19. doi: 10.1016/j.molcel.2008.01.011. PubMed PMID: 18280237.

69. Arigo JT, Eyler DE, Carroll KL, Corden JL. Termination of cryptic unstable transcripts is directed by yeast RNA-binding proteins Nrd1 and Nab3. Mol Cell. 2006;23(6):841–51. Epub 2006/09/16. doi: 10.1016/j.molcel.2006.07.024. PubMed PMID: 16973436.

70. Fasken MB, Laribee RN, Corbett AH. Nab3 facilitates the function of the TRAMP complex in RNA processing via recruitment of Rrp6 independent of Nrd1. PLoS Genet. 2015;11(3):e1005044. Epub 2015/03/17. doi: 10.1371/journal.pgen.1005044. PubMed PMID: 25775092; PubMed Central PMCID: PMCPMC4361618.

71. Lemay JF, D’Amours A, Lemieux C, Lackner DH, St-Sauveur VG, Bahler J, et al. The nuclear poly(A)-binding protein interacts with the exosome to promote synthesis of noncoding small nucleolar RNAs. Mol Cell. 2010;37(1):34–45. Epub 2010/02/05. doi: 10.1016/j.molcel.2009.12.019. PubMed PMID: 20129053.

72. LaCava J, Houseley J, Saveanu C, Petfalski E, Thompson E, Jacquier A, et al. RNA degradation by the exosome is promoted by a nuclear polyadenylation complex. Cell. 2005;121(5):713–24. Epub 2005/06/07. doi: 10.1016/j.cell.2005.04.029. PubMed PMID: 15935758.

73. Shen L, Shao N, Liu X, Nestler E. ngs.plot: Quick mining and visualization of next-generation sequencing data by integrating genomic databases. BMC genomics. 2014;15:284. doi: 10.1186/1471-2164-15-284. PubMed PMID: 24735413; PubMed Central PMCID: PMC4028082.

74. Castelnuovo M, Rahman S, Guffanti E, Infantino V, Stutz F, Zenklusen D. Bimodal expression of PHO84 is modulated by early termination of antisense transcription. Nat Struct Mol Biol. 2013;20(7):851–8. Epub 2013/06/19. doi: 10.1038/nsmb.2598. PubMed PMID: 23770821.

75. Smith-Kinnaman WR, Berna MJ, Hunter GO, True JD, Hsu P, Cabello GI, et al. The interactome of the atypical phosphatase Rtr1 in Saccharomyces cerevisiae. Molecular bioSystems. 2014;10(7):1730–41. doi: 10.1039/c4mb00109e. PubMed PMID: 24671508; PubMed Central PMCID: PMC4074173.

76. Irani S, Yogesha SD, Mayfield J, Zhang M, Zhang Y, Matthews WL, et al. Structure of Saccharomyces cerevisiae Rtr1 reveals an active site for an atypical phosphatase. Sci Signal. 2016;9(417):ra24. Epub 2016/03/05. doi: 10.1126/scisignal.aad4805. PubMed PMID: 26933063.

77. Ni Z, Xu C, Guo X, Hunter GO, Kuznetsova OV, Tempel W, et al. RPRD1A and RPRD1B are human RNA polymerase II C-terminal domain scaffolds for Ser5 dephosphorylation. Nat Struct Mol Biol. 2014;21(8):686–95. Epub 2014/07/07. doi: 10.1038/nsmb.2853. PubMed PMID: 24997600; PubMed Central PMCID: PMCPMC4124035.

78. Chang TK, Lawrence DA, Lu M, Tan J, Harnoss JM, Marsters SA, et al. Coordination between Two Branches of the Unfolded Protein Response Determines Apoptotic Cell Fate. Mol Cell. 2018;71(4):629–36 e5. Epub 2018/08/18. doi: 10.1016/j.molcel.2018.06.038. PubMed PMID: 30118681.

79. Gibney PA, Fries T, Bailer SM, Morano KA. Rtr1 is the Saccharomyces cerevisiae homolog of a novel family of RNA polymerase II-binding proteins. Eukaryot Cell. 2008;7(6):938–48. Epub 2008/04/15. doi: 10.1128/EC.00042-08. PubMed PMID: 18408053; PubMed Central PMCID: PMCPMC2446653.

80. Kapitzky L, Beltrao P, Berens TJ, Gassner N, Zhou C, Wuster A, et al. Cross-species chemogenomic profiling reveals evolutionarily conserved drug mode of action. Mol Syst Biol. 2010;6:451. Epub 2010/12/24. doi: 10.1038/msb.2010.107. PubMed PMID: 21179023; PubMed Central PMCID: PMCPMC3018166.

81. Dronamraju R, Kerschner JL, Peck SA, Hepperla AJ, Adams AT, Hughes KD, et al. Casein Kinase II Phosphorylation of Spt6 Enforces Transcriptional Fidelity by Maintaining Spn1-Spt6 Interaction. Cell Rep. 2018;25(12):3476–89 e5. Epub 2018/12/20. doi: 10.1016/j.celrep.2018.11.089. PubMed PMID: 30566871; PubMed Central PMCID: PMCPMC6347388.

82. Winzeler EA, Shoemaker DD, Astromoff A, Liang H, Anderson K, Andre B, et al. Functional characterization of the S. cerevisiae genome by gene deletion and parallel analysis. Science. 1999;285(5429):901–6. Epub 1999/08/07. PubMed PMID: 10436161.

83. Washburn MP, Wolters D, Yates JR, 3rd. Large-scale analysis of the yeast proteome by multidimensional protein identification technology. Nat Biotechnol. 2001;19(3):242–7. Epub 2001/03/07. doi: 10.1038/8568685686 [pii]. PubMed PMID: 11231557.

84. Mellacheruvu D, Wright Z, Couzens AL, Lambert JP, St-Denis NA, Li T, et al. The CRAPome: a contaminant repository for affinity purification-mass spectrometry data. Nat Methods. 2013;10(8):730–6. Epub 2013/08/08. doi: 10.1038/nmeth.2557. PubMed PMID: 23921808; PubMed Central PMCID: PMCPMC3773500.

85. Knight JDR, Choi H, Gupta GD, Pelletier L, Raught B, Nesvizhskii AI, et al. ProHits-viz: a suite of web tools for visualizing interaction proteomics data. Nat Methods. 2017;14(7):645–6. Epub 2017/07/01. doi: 10.1038/nmeth.4330. PubMed PMID: 28661499; PubMed Central PMCID: PMCPMC5831326.

86. Wickham H. ggplot2 - Elegant Graphics for Data Analysis: Springer-Verlag, New York.; 2016. 260 p.

87. Wal M, Pugh BF. Genome-wide mapping of nucleosome positions in yeast using high-resolution MNase ChIP-Seq. Methods Enzymol. 2012;513:233–50. Epub 2012/08/30. doi: 10.1016/B978-0-12-391938-0.00010-0. PubMed PMID: 22929772; PubMed Central PMCID: PMCPMC4871120.

88. Homer NM, Barry; Nelson, Stanley F. BFAST: an alignment tool for large scale genome resequencing. PLoS One. 2009;4(11):e7767. doi: 10.1371/.

89. Breese MR, Liu Y. NGSUtils: a software suite for analyzing and manipulating next-generation sequencing datasets. Bioinformatics. 2013;29(4):494–6. Epub 2013/01/15. doi: 10.1093/bioinformatics/bts731. PubMed PMID: 23314324; PubMed Central PMCID: PMC3570212.

90. Ching T, Huang S, Garmire LX. Power analysis and sample size estimation for RNA-Seq differential expression. Rna. 2014;20(11):1684–96. doi: 10.1261/rna.046011.114. PubMed PMID: 25246651; PubMed Central PMCID: PMC4201821.

91. Robinson JT, Thorvaldsdottir H, Winckler W, Guttman M, Lander ES, Getz G, et al. Integrative genomics viewer. Nat Biotechnol. 2011;29(1):24–6. Epub 2011/01/12. doi: 10.1038/nbt.1754. PubMed PMID: 21221095; PubMed Central PMCID: PMC3346182.

